# Cyanobacterial cohorts structure the diversity, abundance, and metabolism of heterotrophic bacteria in Lake Erie

**DOI:** 10.64898/2026.08.03.742551

**Authors:** Augustus Pendleton, Sophia Aredas, Kailyn Hanke, Bofan Wei, Gregory L. Boyer, Marian Schmidt

## Abstract

Eutrophication and warming in Lake Erie create two microbial threats: cyanobacterial harmful algal blooms (cHABs) that can produce toxins, and seasonal hypoxia driven by microbial respiration. These phenomena are often studied separately, with cHABs research focused on the Western basin and hypoxia on the Central basin. We conducted lakewide microbial sampling at three time points in 2024 (May, August, and September), integrating physicochemical data, cyanotoxin quantification, amplicon sequencing, and flow cytometry. We demonstrate that cyanobacteria form spatially and seasonally distinct “cohorts” that act as hubs structuring abundant, diverse, and active communities across all basins. These cohorts display distinct relationships with heterotrophic communities, with colonial, bloom-forming cohorts (i.e., *C1: Microcystis*, *C2: Pseudanabaena*) associated with higher richness and evenness, the picocyanobacterium *C3: Cyanobium* with increased heterotrophic abundance, and *C2: Pseudanabaena* additionally associated with increased numbers of metabolically active cells. Neither temperature nor nutrient concentrations consistently explained these patterns, although total phosphorus correlated with bloom-forming C1 and C2 cohorts. The cohorts also formed structured “consortia” with heterotrophic taxa, with each cyanobacterial group associated with consistent sets of heterotrophic partners. Together, these results are consistent with a model in which cyanobacterial abundance increases heterotrophic growth and respiration, suggesting a lakewide pathway linking cHABs to oxygen demand and hypoxia.

**IMPORTANCE:** Cyanobacterial harmful algal blooms and hypoxia are two microbial processes shaping water quality in Lake Erie, yet they are typically studied separately and at basin-specific scales. We link cyanobacterial abundance to heterotrophic metabolism at a lakewide scale. We show that cyanobacterial abundance is associated with heterotrophic diversity, abundance, and metabolic activity, with contrasting patterns across cyanobacterial functional groups. These relationships were not explained by temperature nor consistently by nutrients, although cyanobacterial distribution in the Central basin is likely linked to nutrient availability as a result of upwellings and basin-wide gyres. Bloom-forming and picocyanobacterial cohorts play fundamentally different roles in structuring microbial diversity and biomass, with implications for how bloom management influences ecosystem metabolism. By identifying cyanobacteria as a major predictor of microbial biomass and activity, this work reveals a spatially-explicit pathway connecting cyanobacterial primary production to oxygen demand, suggesting that managing blooms may regulate oxygen depletion.

## INTRODUCTION

Lake Erie supplies drinking water to over 11 million people, yet two of the most consequential threats to its water quality remain mechanistically unlinked: cyanobacterial harmful algal blooms (cHABs) and seasonal hypoxia [1]. Both phenomena have been intensified by eutrophication [2, 3] and warming temperatures [4], yet these processes have largely been studied independently: cHABs are most severe in the shallow, polymictic Western basin, whereas hypoxia occurs primarily in the Central basin during summer stratification. As such, we lack functional predictions between the abundance of cyanobacteria and the activity of associated heterotrophic bacteria on a lakewide scale.

Relative abundance data captures shifts in community composition but does not resolve changes in biomass or metabolic activity [5]. As a result, the extent to which cyanobacterial blooms scale to heterotrophic abundance and respiration remains poorly studied [6, 7, 8]. Absolute quantification is required to link community composition to ecosystem function, particularly for processes such as bloom formation and oxygen consumption that depend on microbial biomass. Liu et al. [9] showed 16S rRNA gene copy abundance did not correlate strongly with a *Microcystis* bloom in Lake Taihu, whereas the metabolic activity (i.e. RNA/DNA ratio) increased, suggesting positive interactions between heterotrophs and cyanobacterial colonies [9].

While studies incorporating absolute abundance are rare, there is substantial work detailing the shifts in bacterial community composition in response to cyanobacteria blooms in other eutrophic lakes and Lake Erie. In Lake Taihu, cyanobacterial abundance correlated to decreases in bacterial diversity and evenness [10]. Work from Lake Erie, however, showed no shifts in richness, but taxonomy-specific responses in evenness [11]. Critically, these studies rely on relative abundance data, limiting the ability to link community structure to biomass and metabolic activity.

Increasing evidence suggests that heterotrophic bacterial activity during blooms is driven by direct metabolic interactions with cyanobacteria rather than parallel responses to environmental conditions alone. The *Microcystis* phycosphere has been linked to nitrogen cycling that can be either competitive (e.g. Qian et al. [12]) or syntrophic (e.g. Li et al. [6]). However, these studies focus primarily on colonial blooms dominated by *Microcystis sp.*, in contrast to the taxonomically diverse filamentous cHABs observed in the Central Basin [13]. Because these interactions depend on the distribution and transport of cyanobacterial biomass, understanding them also requires accounting for the spatial heterogeneity imposed by physical circulation.

Spatial heterogeneity in Lake Erie is shaped by tributary inputs and basin-scale hydrodynamics. The Western basin receives nutrient-rich inflows from the Maumee River, whereas the Detroit River delivers comparatively fewer nutrients, contributing to spatial differences in cyanobacterial bloom intensity [14]. Circulation transports cyanobacterial blooms from the Western basin to the Central basin shorelines [15], where environmental conditions differ between the northern and southern shores. The northern shore experiences higher erosion [16] and frequent upwelling events that can increase nutrient availability [17]. Under prevailing winds, circulation in the Central basin is clockwise, generating coastal jets that redistribute water masses west-to-east along the northern shore [18, 19]. How hydrodynamic processes shape (cyano)bacterial communities and mediate their transport between basins remains unresolved, as defining these interactions requires microbial sampling at a broad spatial scale [20].

We hypothesize that cyanobacterial cohorts structure heterotrophic bacterial composition, abundance, and metabolism. We define “cohorts” as recurring, spatially structured assemblages of co-occurring cyanobacterial taxa, and refer to the associated heterotrophic communities they organize as “consortia.” Under this framework, we expect (i) taxonomically structured co-occurrence between cyanobacteria and heterotrophs, (ii) positive correlations between heterotrophic abundance and metabolic activity with cyanobacterial abundance, and (iii) these relationships reflect taxonomically specific associations between cyanobacterial cohorts and heterotrophic consortia, rather than a uniform response to shared physicochemical conditions.

To test this hypothesis, we combine lakewide sampling of Lake Erie with absolute quantification of microbial abundance and activity. We identify spatially and seasonally distinct cyanobacterial cohorts and evaluate the heterotrophic consortia they organize, including their taxonomic composition and their relationship to heterotrophic abundance and metabolic activity. By integrating quantitative, taxonomic, and functional data across the lake, we assess whether cyanobacterial productivity increases heterotrophic respiration independently of environmental co-variates. Our data support a model in which either Maumee-derived (in the Western Basin) or upwelling-derived (in the Central Basin) nutrients foster cyanobacterial blooms, which in turn support highly-diverse, metabolically-active heterotrophic consortia, pointing to a mechanistic connection between blooms, microbial diversity, and oxygen demand.

## MATERIALS AND METHODS

Methods are condensed for brevity, but full experimental details are available in the Supplemental Information.

### Sample collection

Samples were collected in 2024 in collaboration with the US Environmental Protection Agency (EPA) aboard the *R/V Lake Guardian* (Fig. 1). A total of 33 stations were visited at each cruise; complete sampling dates and locations can be found in Table S1. At all stations, depth profiles were collected via a conductivity, temperature, and depth (CTD) meter.

**FIG 1.**
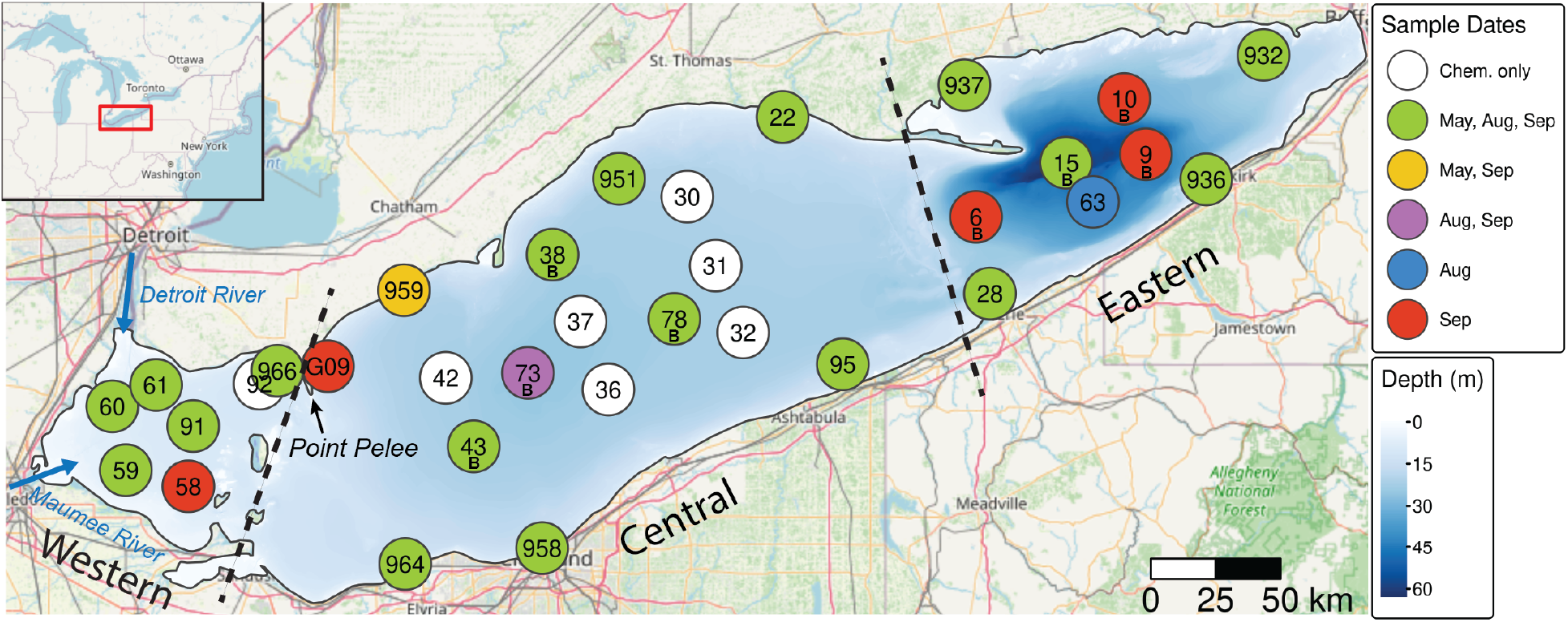
Sampling Locations and Dates in Lake Erie. ”Integrated” surface samples (see Methods, “Sample Collection”) were collected for microbial analysis from 26 stations in May, August, and September of 2024, though not all stations were sampled each month (symbol color; no microbial samples were collected at white stations). At all stations, the water column was profiled via a Conductivity, Temperature, and Depth (CTD) cast with additional sensors, as well as sampled for comprehensive EPA water quality testing. Stations with a corresponding “B” also had bottom samples collected 1m above the lakebed. Lake Erie’s three basins (Western, Central, and Eastern) are demarcated with dashed lines.

Depth-discrete water samples for environmental data were collected using an attached Niskin-bottle rosette, with “surface” samples taken at 2 m deep; when stations were sufficiently deep to observe a thermocline, additional “bottom” samples were collected 2 m above the lakebed. Chemical parameters and chlorophyll-a concentrations were analyzed by the US EPA’s Great Lakes National Program Office using standard protocols [21].

Microbial sampling was carried out at as many stations as possible, given time constraints of shipboard sampling (Fig. 1). At each station, an integrated epilimnetic sample was generated by combining equal volumes of water from 1 m, 5 m, and 10 m depths, hereafter referred as “surface” samples. This sampling approach was used because bloom-forming cyanobacteria like *Microcystis sp.* can be heterogeneous within the epilimnion [22]. For amplicon sequencing, 2 L of water was filtered through a 0.22 *µ*m polyethersulfone (PES) filter (MilliporeSigma) and then flash frozen in liquid N_2_ before storage at -80°C. Filtration lines were sterilized with bleach and MilliQ water between samples. For particulate toxin analysis, 2.5 L of water was filtered through 0.7*µ*m GF/F glass fiber filters (Whatman) and samples were stored at -20°C. Between toxin samples, filtration lines were rinsed only with MilliQ water. At the end of each cruise, field negative controls for both DNA and toxin filters were produced by running 2L of MilliQ onto either 0.22 *µ*m PES or 0.7*µ*m GF/F filters after standard cleaning procedures.

Samples for flow cytometry were collected from integrated eplimnetic samples prefiltered through sterile 200*µ*m and 20 *µ*m Nitex mesh (Wildco) to prevent clogging. All counts were performed using an Attune Nxt flow cytometer (Thermofisher) equipped with a small-particle filter. For metabolic labeling with RSG, which was only collected in September, 1 mL of prefiltered sample was first incubated with 1 *µ*L of 1mM RSG (Thermofisher) for 30 minutes in the dark at room temperature, after which 1*µ*L of 25% glutaraldehyde was added and samples were incubated for an additional 10 minutes. All samples were collected in biological duplicates, flash frozen in liquid N_2_, and stored at -80°C.

### Cyanotoxin Quantification

Filters for cyanotoxin analysis were transferred to SUNY-ESF for analysis. They were extracted in 50% acidified methanol using ultrasound, centrifuged at 14,000 x g, and filtered through a 0.22 *µ*m nylon filter prior to storage at -20°C. Microcystins (17 congeners) were analyzed by HPLC-coupled with single quadrupole mass spectrometry and quantified against a standard curve of microcystin LR or RR as outlined in Boyer [23]. Anatoxin-a, homo-anatoxin-a, cylindrospermopsin, and deoxy-cylindrospermopsin concentrations were determined using LC-MS/MS as described in Smith et al. [24]. The method detection limits for all toxins were <0.01 *µ*g/L.

### Cell quantification via flow cytometry

Flow cytometry was used to quantify three microbial properties: (1) total microbial cell abundances, (2) phototrophic cells based on chlorophyll-a autofluorescence, and (3) metabolically active cells using RedoxSensor Green (RSG) labeling (Fig. S1). Total cells were quantified based on DNA staining with SYBR Green I [25]. Phototrophic cells were quantified based on chlorophyll-a autofluorescence [26]. Cell size was estimated using the forward scatter (FSC-H) after creating a standard curve using a Flow Cytometry Size Calibration Kit (Thermofisher; F13838), with a 5*µ*m cutoff for “Large” cells (Fig. S1). Metabolically active cells were quantified using RedoxSensor Green (RSG) [27]. Full details of flow cytometry staining, collection parameters, laser voltages, thresholds, and gating are available within Supplemental Methods and Table S2.

### DNA Extraction and Metabarcoding

All DNA extractions were carried out using the Qiagen DNeasy PowerWater kit per manufacturer’s protocol. The V4–V5 hypervariable region was amplified using the universal primers 515F (5-GTGYCAGCMGCCGCGGTAA) and 926R (5-CCGYCAATTYMTTTRAGTTT) as described in Yeh et al. [28] and Needham et al. [29]. In total, 80 microbial samples were sequenced along with negative controls (four field blanks, two extraction blanks, one PCR blank, and one indexing blank) and a defined microbial community (ZymoBIOMICs Microbial Community DNA Standard) used as a positive control to assess amplification error rates. Libraries were sequenced at the Cornell Biotechnology Resources Center using a 2 x 300 bp paired-end XLEAP kit on an Illumina NextSeq 2000.

Sequencing data were processed to generate amplicon sequence variants (ASVs) from prokaryotic reads using scripts modified from McNichol et al. [30]. Sequences were denoised and ASVs inferred using a standard DADA2 workflow, with minor adjustments for binned quality scores [31].

ASVs were taxonomically classified using the Greengenes2 database (v2024.09) [32]. Mitochondrial and chloroplast ASVs were removed, alongside two additional ASVs that were present in the negative controls or appeared to originate from mock community contamination. To reduce potential erroneous ASVs, a minimum observation threshold of eight reads per ASV was applied, leaving 10,774 ASVs but retaining 99.9% of all sequencing reads.

A phylogenetic tree was constructed using MAFFT for sequence alignment and FastTree for tree inference under a generalized time-reversible model [33, 34]. Archaea were poorly represented (three ASVs); given previous documentation of archaeal taxa in Lake Erie, this scarcity likely reflects low amplification efficiency for Archaea with these primers despite positive *in silico* predictions [30, 35]. Samples with less than 10,000 reads were removed. One sample (Stn. 937 in September) exceeded this threshold (14,676) but exhibited anomalously low richness and cell counts, suggesting possible residual bleach contamination in the sampling bottle, and was therefore removed. Of the original 80 samples sequenced, nine were excluded from final analyses due to insufficient sequencing depth or quality (Table S3).

### Ecological analyses

Alpha-diversity (richness) was estimated after rarefying samples to the minimum sequencing read depth (11,901 reads) [36].

Beta-diversity was calculated using Bray-Curtis dissimilarity based on absolute abundances [37]. Because flow cytometry counts were generated from the <20 *µ*m fraction whereas sequencing was performed without any prefiltration, colonial cyanobacteria were likely underestimated during flow cytometry and compared to their total relative abundance in sequencing reads. Cyanobacterial spatial groups were defined using UPGMA clustering, cutting the tree at 7 groups to maximize clustering by both month and basin.

Co-occurrence network analyses were performed using functions from the igraph and tidygraph packages [38, 39]. For cyanobacteria–cyanobacteria co-occurrence, the 100 most abundant cyanobacterial ASVs were selected with a prevalence of at least 25%. Pairwise associations were assessed using Pearson correlations, and significant correlations (edges) were defined as r *≥* 0.8 and p *≤* 0.01 (uncorrected). These edges were used to construct a weighted, undirected network that maximized modularity (Fig. S2). For the cyanobacteria–heterotroph co-occurrence network, a minimum absolute abundance of > 50,000 cells/ml and 25% prevalence was applied, leaving 344 heterotrophic ASVs (cyanobacterial ASVs were removed, see Supplemental methods, Fig. S2 and Fig. S3). Pearson correlations were then calculated between each of the 100 cyanobacterial ASVs (”hubs”) and heterotroph ASVs (”nodes”). Edges were defined as r *≥* 0.85 and p < 0.01 (uncorrected) and used to construct a weighted, undirected bipartite network. For both networks, clusters were identified using the Givan-Newman algorithm and networks were visualized using the Fruchterman-Rheingold layout.

General data manipulations relied on the tidyverse and phyloseq packages using R v4.3.3 [40, 41, 42]. Other statistical tests (including Two-Sample Wilcoxon Tests and Spearman Correlations) were performed using functions from base R, rstatix, or vegan packages [43, 44]. Plots were produced using functions from the ggplot2, ggpubr, and ggdendro packages [41, 45, 46].

## RESULTS

### Bacterial and phototrophic abundances vary across basin-level environmental gradients

Total and phototrophic cell abundances were lowest in May and increased during summer sampling (August and September), consistent with seasonal bloom development (Fig. S4A-C). Total cell abundances varied across basins but did not follow the west-to-east decline predicted by general water and nutrient transport across Lake Erie. The Western basin was the most heterogeneous, with high abundances at southern stations and low abundances at northern stations (Fig. 2A), consistent with elevated nitrogen and phosphorus from the Maumee River relative to the Detroit River (Fig. S5). Abundances in the Central basin were comparable to those in the Western basin and significantly higher than the Eastern basin (Fig. S4A), but exhibited a reversed spatial structure with higher cell abundances along the northern than the southern shore (Fig. 2A). Central and Eastern basin abundances were vertically structured, with reduced cell counts in the hypolimnion (Figs. 2A, S4D, G). In the Central basin, this pattern coincided with widespread hypoxia in August and September (Fig. S6).

**FIG 2.**
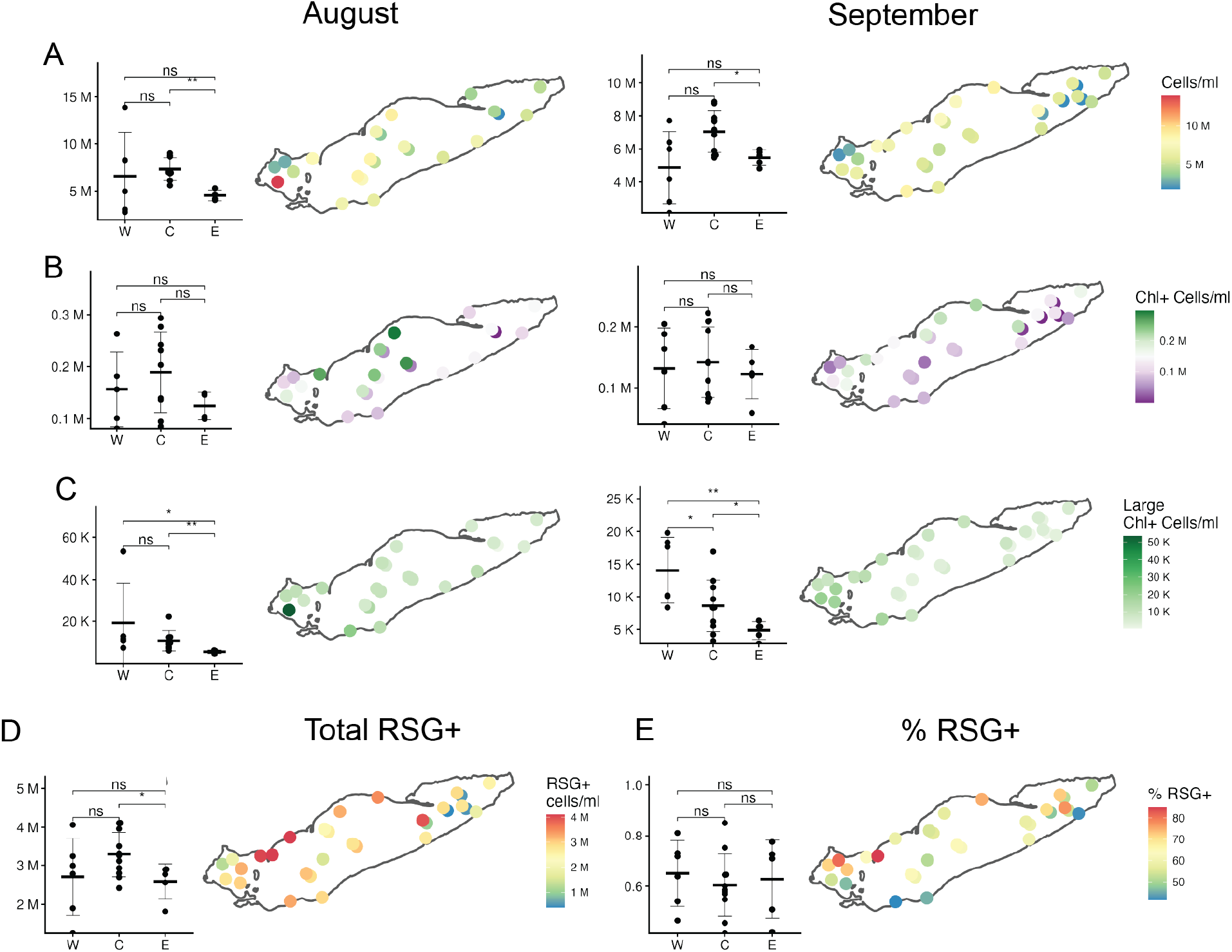
Bacterial and cyanobacterial abundances are spatially structured across the three basins of Lake Erie. (A) Total surface bacterial cell abundance, (B) chlorophyll-a + surface cells, (C) large (>5*µ*m) chlorophyll-a + cells (D) metabolically active or RSG+ cells (September only), and (E) the proportion of RSG+ cells (September only) measured by flow cytometry after <20*µ*m prefiltration. Inset plots show surface samples grouped by basin (Western (W), Central (C), and Eastern (E)). Error bars reflect the mean *±* standard deviation. Stars represent results of Two-Sample Wilcoxon tests with Holm-Bonferroni adjustment for multiple comparisons (* = p < 0.05, ** = p < 0.01, *** = p < 0.001, **** = p < 0.0001). Points represent surface samples, with bottom samples for those stations plotted offset underneath.

Phototrophic cells exhibited similar spatial patterns to total cell abundances, but with significant intra-basin variability (Figs. 2B-C). In the Western basin, Chl+ and large Chl+ cells were most abundant in the south and reduced at northern stations. In the Central basin, this pattern was reversed, with higher phototroph abundances along the northern than the southern shore. Current models in the week before and during sampling support upwelling along the Central basin’s north shore and water movement into the offshore, potentially stimulating cyanobacterial growth in the nearshore and transporting cyanobacteria from the nearshore into the offshore (Fig. 3). Large Chl+ cells, which likely correspond to colonial or filamentous species associated with HABs, decreased in abundance from the Western to the Eastern basin, especially in September (Fig. 2C). Phototroph abundances were lowest and more spatially uniform in the Eastern basin.

**FIG 3.**
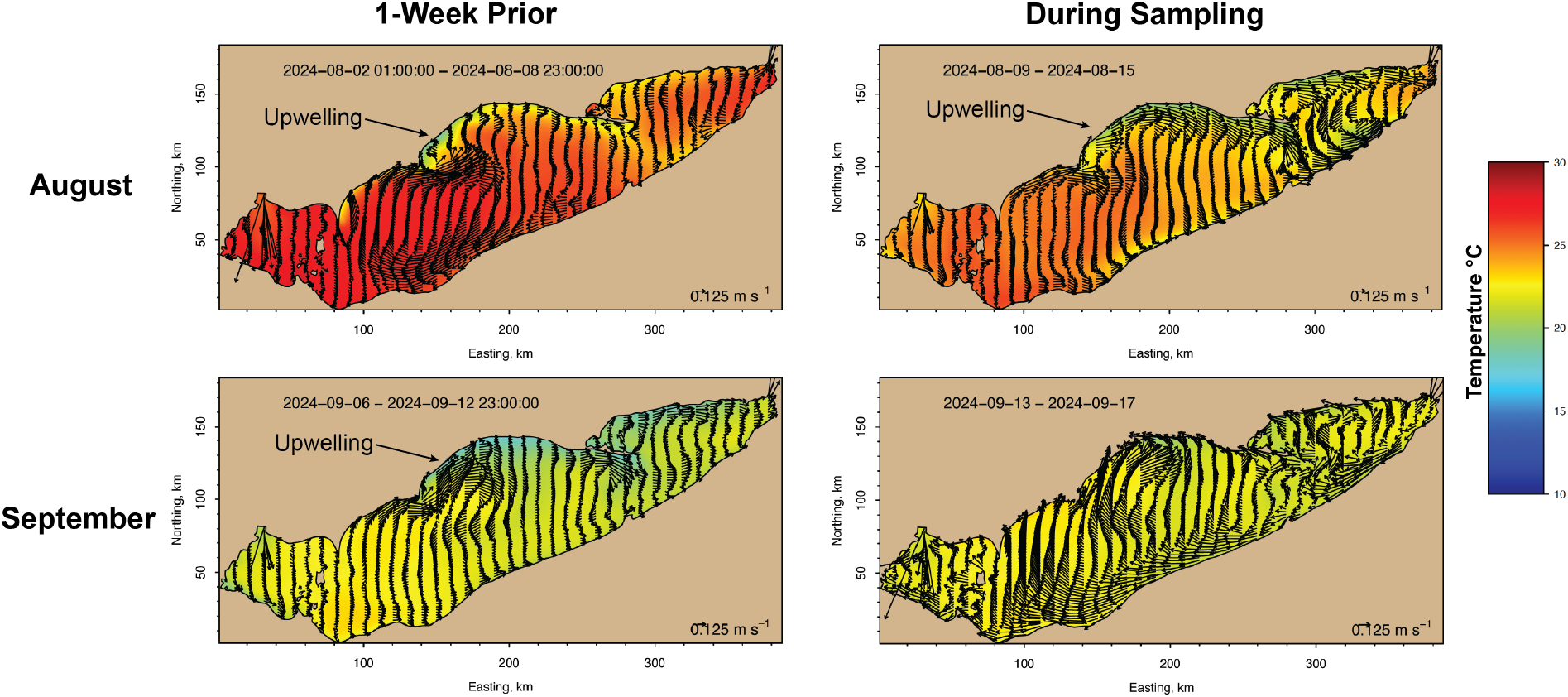
Surface temperature and current models before and during sampling. Data was accessed from the NOAA Lake Erie Operational Forecasting System (LEOFS) FVCOM model, using the n000 nowcast results.

Metabolically active (RSG+) cells were only measured in September and displayed clear spatial structure. RSG+ cell abundances were moderate in the Western basin, highest in the Central basin, and lowest in the Eastern basin. Within the Central basin, peak abundances occurred along the northern shore (Fig. 2D). In contrast, the proportion of active cells (%RSG+) ranged from 42% - 85% across all three basins and did not follow the same spatial patterns, with the highest values close to Point Pelee (Fig. 2E). Consistent with this, %RSG+ had no clear relationship with total cell abundances (Spearman’s Rank Correlation = 0.0031, p = 0.993). Notably, high-abundance bloom samples in the southern Western basin did not exhibit elevated %RSG+.

### Toxin-producing cyanobacteria form spatially structured cohorts

Particulate cyanotoxins showed strong spatial patterns and taxon-specific associations across the lake (Fig. 4). Microcystins were localized to the Western basin and peaked in August (0.29 *µ*g/L, 75% LR/25% YR, Fig. 4A), coinciding with elevated absolute abundances of *Microcystis* (Fig. 4C). In contrast, anatoxins were detected at lower concentrations (maximum 0.048 *µ*g/L) and were concentrated in the Central basin, especially near Point Pelee in September (Fig. 4B). Anatoxin concentrations were most strongly associated with the absolute abundance of *Planktothrix sp.* (Figs. 3D, S7). Cylindrospermopsins were not detected.

**FIG 4.**
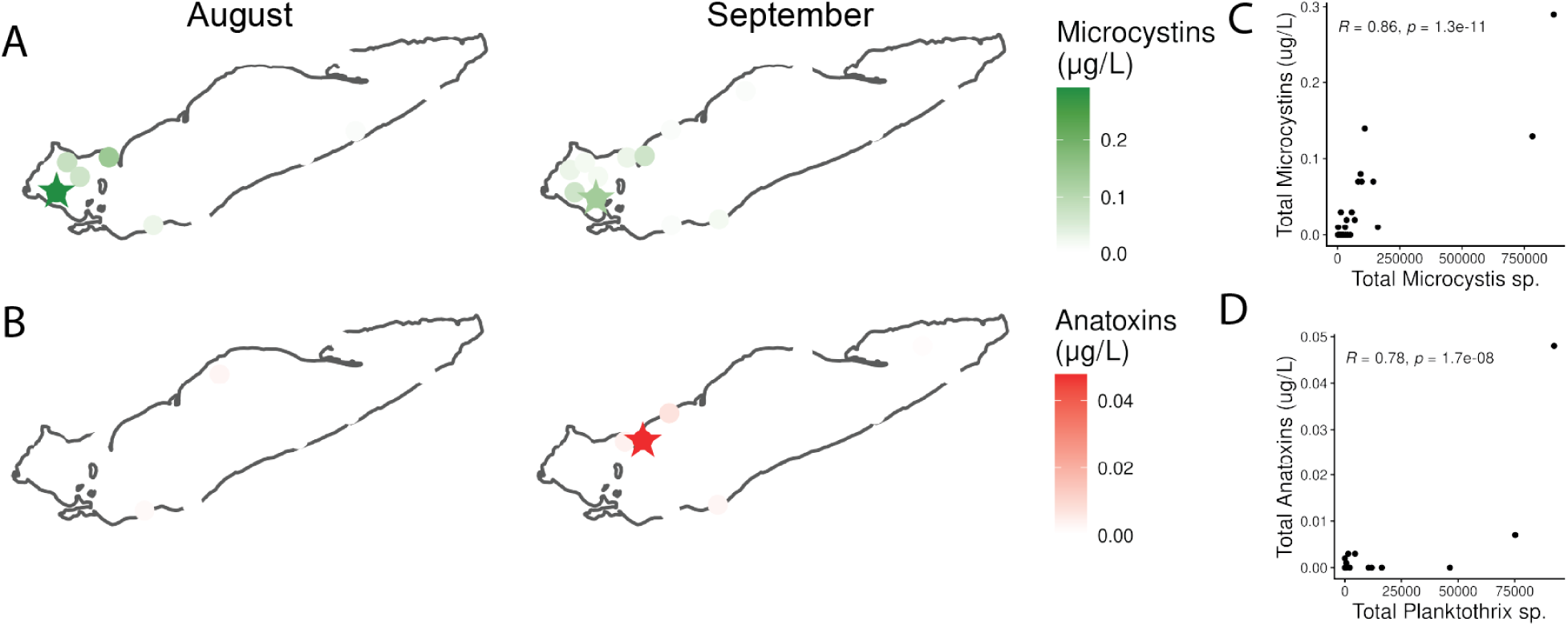
Cyanotoxins distributions track spatial dominance of distinct cyanobacterial genera. (A) Total particulate microcystins and (B) total particulate anatoxins. (C) Total microcystin content is plotted versus absolute counts of ASVs in genus *Microcystis*. (D) Total anatoxin content is plotted versus absolute counts of ASVs in genus *Planktothrix.* R and p-values represent Spearman’s Rank Correlation.

Cyanobacterial communities were spatially structured across Lake Erie’s surface waters, forming seven clusters aligned with basin and community composition (Fig. 5). Western basin samples were dominated by *Microcystis*, with “Peak Bloom” clusters characterized by high absolute abundances. “Bloom Adjacent 2” communities retained high relative abundances of Microcystaceae but lower total cell abundances. In contrast, Central basin communities differed between months and were taxonomically diverse with high absolute abundances. They shared a common backbone of *Cyanobium*, Microcystaceae, Microcoleaceae, *Dolichospermum*, and *Pseudanabaena*, with multiple filamentous lineages present in both months but differing in composition (e.g. *Limnoraphis* in August and *Nodosilinea* in September). Abundances shifted from relatively even distributions in August to increased dominance by Microcystaceae-, *Dolichospermum*-, and *Pseudanabaena*-associated lineages in September. Eastern basin communities in both months were dominated by the picocyanobacteria *Cyanobium*, with moderate cell counts and minimal representation of bloom-forming taxa. Two samples (St. 15 in August, “Deep Desert”, and St. 60 in September, “Bloom Adjacent 1”) had exceptionally low cyanobacterial abundance.

**FIG 5.**
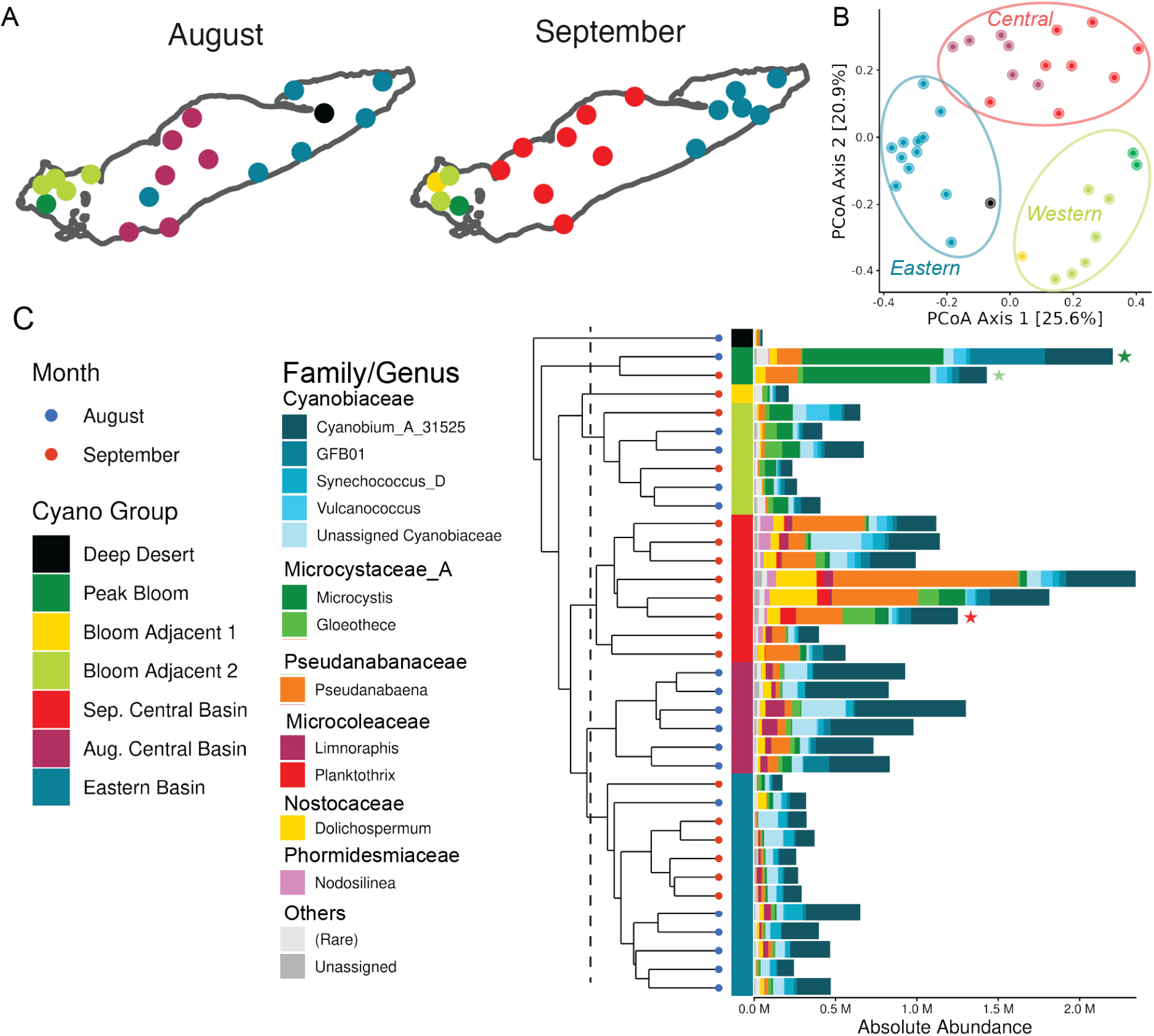
Cyanobacterial communities form spatially structured assemblages defined by abundance and composition. (A) UPGMA clustering of cyanobacterial communities in August and September based on the absolute abundance-weighted Bray-Curtis dissimilarity. The underlying dendrogram is displayed in (C), with the dashed line corresponding to cut height resulting in seven clusters. (B) Principal Coordinates Analysis, also based on the absolute abundance-weighted Bray-Curtis dissimilarity, demonstrating group clustering. (C) Cyanobacterial cell abundance and community composition across these clusters, arranged at the Genus level. Samples with highest toxin levels from Fig. 4 are annotated with corresponding stars.

Abundant cyanobacterial ASVs formed three major co-occurrence clusters (”cohorts”) with distinct spatial and seasonal distributions (Fig. 6). The first cohort (”*C1: Microcystis*”) resembled “Peak Bloom” communities and was dominated by *Microcystis*, with strong connectivity between *Microcystis* and ASVs within the Greengenes2 genus *GFB01* (Family Cyanobiaceae). A second cohort (”*C2: Pseudanabaena*”) was associated with the highly diverse cyanobacterial assemblages observed in the Central basin in September and included taxa such as *Pseudanabaena*, *Dolichospermum*, and *Planktothrix*, which are known for colonial lifestyles and cyanotoxin production. The third cohort (”*C3: Cyanobium*”) was dominated by *Cyanobium sp.* and occurred throughout the Central basin in August and along its northern shore in September.

**FIG 6.**
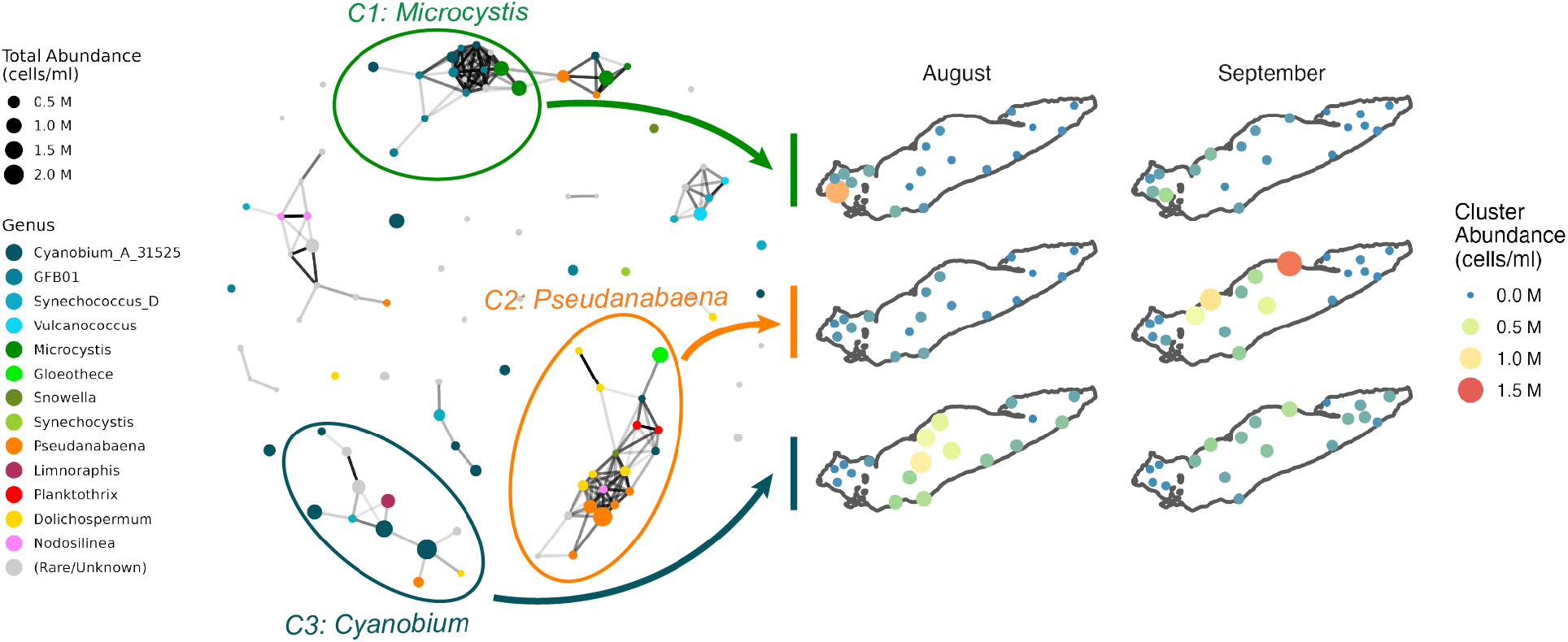
Cyanobacteria form taxonomically consistent co-occurrence “cohorts”. The 100 most abundant cyanobacterial ASVs in August and September were selected for co-occurrence analysis, with a prevalence cut-off of at least 25%. The Pearson correlation was calculated between all ASV pairs, an adjacency matrix was created with *r* > 0.8, p < 0.01, and the network was visualized via the Fruchterman-Reingold algorithm. Nodes (ASVs) are colored by Genus, and their size corresponds to the sum of their absolute abundance across all samples. Clusters (large ellipses) were identified via the Givan-Newman algorithm. The summed abundance of all ASVs in each cluster (”Cluster Abundance”) within each sample was then plotted in August and September. Both the color and size of map points correspond to the cluster abundance, not taxonomic identity.

### Cyanobacterial cohorts support abundant, metabolically active heterotrophic consortia

We sought to link these discrete cyanobacterial assemblages to the heterotrophic bacteria associated with them. Heterotrophic richness and evenness were not correlated with total cyanobacterial abundance but were positively associated with the colonial cohorts *C1: Microcystis* and (evenness only) *C2: Pseudanabaena*. In contrast, total heterotrophic abundance increased with cyanobacterial abundance and was most strongly associated with *C3: Cyanobium* (Fig. 7, Fig. S8).

**FIG 7.**
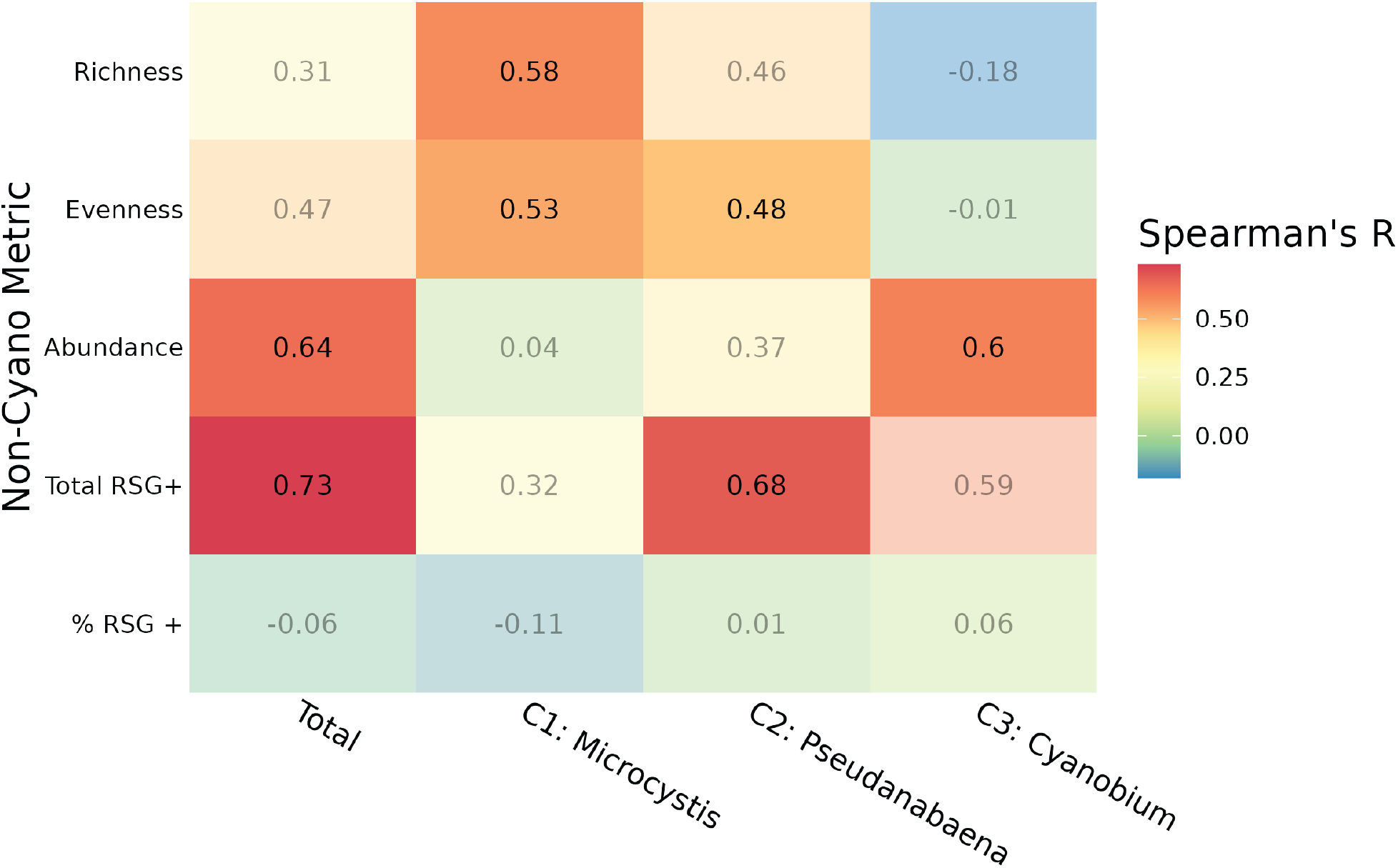
Cyanobacterial abundance and cohort identity correlate with non-cyanobacterial richness, abundance, and metabolic activity. Spearman Rank Correlations (R) were calculated between the summed abundance of cyanobacterial cohorts and heterotrophic metrics. “Total” corresponds to all cyanobacteria, while C1–C3 correspond to the three cyanobacterial cohorts identified in Fig. 6 (*Microcystis*, *Pseudanabaena*, and *Cyanobium*). Heterotrophic metrics (excluding cyanobacterial ASVs and their counts) include richness, evenness, and total cell abundance, the number of RSG+ (metabolically active) cells, and the proportion of RSG+ cells in the community (%RSG+). Color indicates the Spearman rank correlation coefficient, with values shown in each panel. Transparent panels were non-significant relationships (p > 0.05 with a Holm-Bonferroni adjustment). RSG measurements were only available for September samples. Corresponding scatterplots are shown in Fig. S8.

These relationships were not explained by temperature or nutrients (Figs. S9, S10). None of the heterotrophic community metrics were correlated with temperature (Fig. S9), and no consistent relationships were observed with total nitrogen (TN), total phosphorus (TP), or N:P ratios (Fig. S10). Although heterotrophic richness and evenness were positively correlated with TP (Fig. S10), TP also correlated with the abundance of the colonial, bloom-forming cyanobacterial cohorts (*C1: Microcystis/C2: Pseudanabaena*, R = 0.7, p < 0.001), but not with the picocyanobacterial *C3: Cyanobium* cohort (Figs. S9B–C).

Absolute metabolic activity of heterotrophic cells (RSG+ cells mL*^−^*^1^, excluding Chl+ cells) increased with cyanobacterial abundance and was positively associated with the *C2: Pseudanabaena* cohort (Fig. 7). However, the proportion of metabolically active cells (% RSG+ within the community, excluding Chl+ cells) did not correlate with cyanobacterial abundance (Fig. 7).

To assess what specific interactions may drive these community-wide linkages, we tested co-occurrence relationships between cyanobacterial and heterotrophic ASVs. Abundant cyanobacterial cohorts formed structured co-occurrence networks with heterotrophic ASVs, resolving into three major “consortia” (Fig. 8). Each consortium was organized around a dominant cyanobacterial “hub” (triangles) and included distinct sets of heterotrophic “nodes” (circles). A *Synechococcus*-associated consortium (light blue triangle) included members of the Bacteroidota (yellow circles: *Algoriphagus*, *Dinhuibacter*, and *TMED14*) and Actinomycetota (red circles: *Nanopelagicus* and *BACL27*). A *Cyanobium*-associated consortium (dark blue triangles) associated with *Sediminibacterium*, which also co-occurred with *Limnoraphis*. In contrast, a *Pseudanabaena*-*Dolichospermum*-associated consortium (orange and yellow triangles) included more taxonomically diverse taxa and poorly resolved partners, including Chloroflexota (Caldilineaceae) and members of the Phyla Patescibacteria, Acidobacteriota, Bacteroidota, and Planctomycetota (Genus *Fuerstiella*). Notably, *Microcystis* ASVs had limited connectivity, with a single strong association to an Actinobacteriota ASV within the *Luna1* group [47].

**FIG 8.**
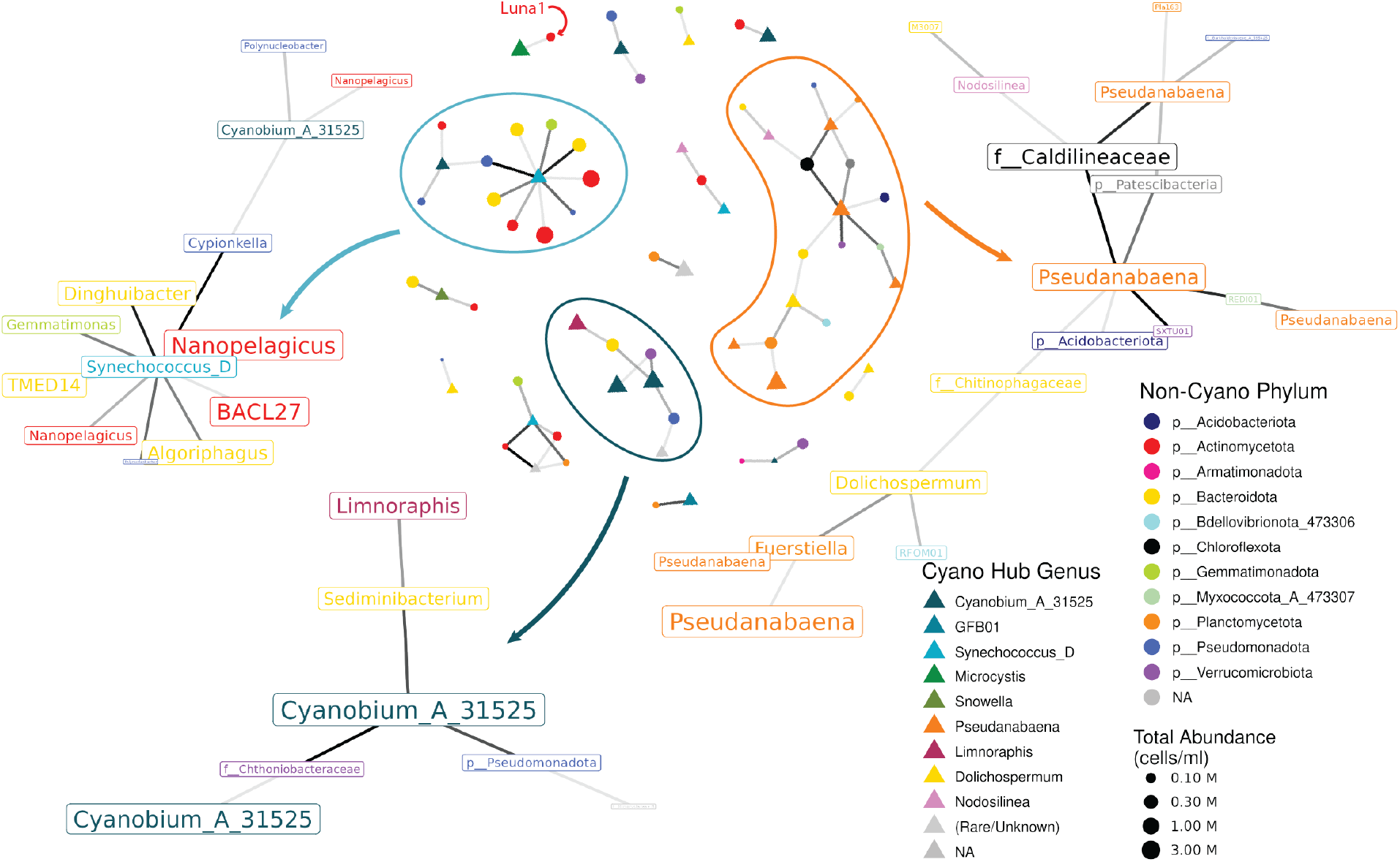
Cyanobacterial cohorts form taxonomically diverse consortia with heterotrophic bacteria. Non-cyanobacterial ASVs with a total abundance > 50,000 cells/ml and prevalence >25% (August and September) were selected for co-occurrence analysis with the 100 most abundant cyanobacterial ASVs (Fig. 6). Pairwise Pearson correlations were calculated between ASVs, and a bipartite network was constructed linking cyanobacterial “hubs” (triangles, colored by Genus) to co-occurring heterotrophic ASVs (circles, colored by Phylum). Both circles and triangles are sized by total abundance of the ASV they represent. Edges represent strong positive associations (R *≥* 0.85, p < 0.01). The three largest clusters, identified via the Givan-Newman algorithm are shown. Genus-level classifications are provided where available.

## DISCUSSION

By integrating quantitative, taxonomic, and metabolic data at a lakewide scale, our results are consistent with cyanobacterial abundance increasing heterotrophic biomass and respiration. Cyanobacteria formed spatially-defined cohorts, which in turn structured taxonomically-specific heterotrophic consortia. The interaction between these cohorts and their consortia, however, depended on cyanobacterial cohort identity: the C1: *Microcystis* cohort correlated to higher richness and evenness; the C2: *Pseudanabaena* cohort correlated to higher evenness and metabolically-active cells, and the picocyanobacterial cohort (C3: *Cyanobium*) was associated with increases in total heterotrophic abundance.

### Cyanobacterial cohort identity structures heterotrophic consortia

Previous studies have demonstrated the cyanobacterial phycosphere creates distinct heterotrophic communities which interact with their algal hosts, though with a primary focus on *Microcystis*. Berry et al. [11] found that heterotrophic evenness generally increased during *Microcystis* blooms. Heterotrophic communities are specific to *Microcystis* oligotype [48, 49], colony size [50], and bloom stage [11], and carbon, nitrogen, and vitamin exchange has been demonstrated between *Microcystis* and its consortia [6, 7].

While these studies remain relevant to the Western Basin where *Microcystis* dominates, the Central basin hosts highly abundant, diverse assemblages of filamentous cyanobacteria (e.g. *Pseudanabaena*, *Dolichospermum*, and *Planktothrix*) which also supported higher evenness. There is evidence of distinct phycosphere communities for these other species (e.g. Fuster et al. [51], Yan et al. [52], Louati et al. [53]), though direct metabolic exchange has not been demonstrated to our knowledge. Limited studies of biotic interactions between heterotrophs and freshwater picocyanobacteria demonstrate tight taxonomic coupling, though to our knowledge direct nutrient exchange within these consortia has not been demonstrated [54].

### Cyanobacterial abundance increases community-level respiration potential

The Central basin cohorts supported higher absolute numbers of RSG+ cells, suggesting cyanobacterial productivity encourages microbial respiration. Despite this, the percentage of the heterotrophic community which was RSG+ was not correlated to cyanobacterial abundance. Munson-McGee et al. [27] showed the most abundant lineages of bacterioplankton in the pelagic ocean had low RSG staining, potentially reflecting rhodopsins supporting ATP-synthesis outside of respiration. If true in Lake Erie, it would suggest that cyanobacteria support the abundance of these lineages not through direct feeding of carbon, but more subtle interactions such as vitamin or nutrient exchange [55, 56]. These targeted interactions may also support the taxonomic specificity and co-occurrence relationships which we observed [57].

### Cyanobacteria-heterotroph consortia are taxonomically specific

Two of our largest consortia centered around picocyanobacteria which were widespread throughout Lake Erie (*Synechococcus* and *Cyanobium*). The first co-occurrence cluster, centered around *Synechococcus_D*, featured highly abundant ASVs in the Actinomycetota alongside ASVs in the Bacteroidota (*Algoriphagus*). Direct physical associations between the polysaccharide-degrading *Algoriphagus sp.* and *Synechococcus sp.* in benthic biofilms have been demonstrated in saltwater environments, and co-occurrence has been widely reported in freshwater [58, 59, 60], emphasizing the potential importance of this relationship.

Highly abundant Actinomycetota, including *Nanopelagicus* and *BACL27* were also linked to *Synechococcus_D* (Fig. 8). The cosmopolitan, highly abundant lineages in the Actinomycetota (e.g. *acI, acIV, Luna*) have also demonstrated niche differentiation which is responsive to *Microcystis* blooms [11, 61]. Intriguingly, Berry et al. [11] noted high co-occurrence of Synechococcus during *Microcystis* blooms, and in our own dataset, a closely related genus, *GFB01*, strongly co-occurred within the C1: *Microcystis* cohort. Only *Luna1* was linked to *Microcystis*, complicating whether these *Actinomycetota* are responding to *Microcystis* or instead commonly co-occurring picocyanobacteria.

The second picocyanobacteria co-occurrence cluster centered around the most dominant ASVs in the genus *Cyanobium*. This cluster recapitulated previous interactions between *Sediminibacterium* and *Cyanobium* but suggested additional partnerships between *Sediminibacterium* and *Limnoraphis* [62]. Picocyanobacteria (C3: *Cyanobium*) also drove positive correlations between cyanobacterial abundance and heterotrophic abundance (Fig. 7). This emphasizes the importance of picocyanobacterial abundance not just as primary producers but determinants of the taxonomy and abundance of heterotrophic bacteria throughout the water column.

The third consortium was anchored by *Pseudanabaena* and *Dolichospermum*, and featured taxonomically diverse heterotrophic partners, in concordance with the C2: *Pseudanabaena* cluster supporting heterotrophic evenness. Many of these partners had poor taxonomic identification, including ASVs in the Phyla Acidobacteriota and Patescibacteria, abundant ASVs in the Families Caldilineaceae (Chloroflexota) and Chitinophagaceae (Bacteroidota), and in the genus *Fuerstiella* (Planctomycetota). The co-occurrence of relatively rare ASVs with these filamentous bacteria in conjunction with their correlation to evenness may imply that more diverse cyanobacterial blooms in the Central basin in turn support a more diverse (and metabolically active) heterotrophic consortium featuring rare taxa, as originally proposed by Berry et al. [11]. While associations between Chitinophagales (and other polysaccharide-degraders) and *Dolichospermum* have been identified previously [51, 53] it is difficult to predict what specific interactions may be occurring for other heterotrophic partners without better taxonomic resolution or metagenomic/metatranscriptomic sequencing.

### Lakewide physical processes structure biological interactions

The biological interactions we predict, while playing out on the scale of microns, are structured by physical processes occurring on a lakewide scale. Heterotrophic community metrics were not explained by temperature and were not consistently associated with nutrient concentrations, although total phosphorus strongly covaried with bloom-forming cohorts. This pattern suggests that nutrients influence cyanobacterial abundance, which in turn structure heterotrophic communities through biological interactions, though we cannot fully exclude TP as an independent driver of heterotrophic richness and evenness.

In the Central basin, we identified linkages between upwelling along the northern shore and cyanobacterial abundance and diversity (Fig. 2, Fig. 3). The enrichment of nutrients (including phosphorus, nitrogen, and iron) within hypoxic bottom waters and their subsequent movement to the surface during upwellings have been shown in Lake Erie [63, 64]. These nutrient pulses have been linked to cyanobacterial abundance along the northern shore [65], and enrichment of soluble active phosphorus specifically in the growth of *Pseudanabaena* and *Planktothrix* over *Microcystis* as we observe here [66].

These dynamics are critical given that *Planktothrix* was highly correlated to anatoxins in our samples, and the low N:P conditions created by upwellings could in turn stimulate toxin production [67]. Anatoxin production by *Planktothrix* has been reported (e.g. Viaggiu et al. [68]), but noy yet from Lake Erie, and our correlational data requires confirmation in culture. The rotation of the gyre could also be moving *Planktothrix* from Sandusky Bay to the northern shore, though we are unable to test this wtihout data from outside the Sandusky outlet. Since the early 2000s, microcystin-producing *Planktothrix* blooms were common in Sandusky Bay, though have been declining since 2019 [69]. Altogether, the surprising appearance of *Planktothrix* alongside anatoxins on the northern shore means monitoring in these regions is essential, in addition to active monitoring in the Western Basin and Sandusky Bay.

The prevailing movement of the gyre in the Central Basin moves both water and microbes east along the northern shore, and then out into the center of the Lake [18, 17]. The movement of nutrients and the cyanobacterial productivity associated with them into the pelagic offshore of Lake Erie due to the prevailing gyre has not been demonstrated to our knowledge, despite widespread pelagic cyanobacterial abundance [13]. This exact hypothesis has been proposed for Lake Biwa, where cyanobacterial biomass is generated in the nutrient-rich nearshore and concentrated into the center of the pelagic gyre [70]. This hydrodynamic link could play a key role in encouraging both cyanobacterial productivity and associated heterotrophic respiration in the Central basin, intensifying hypolimnetic oxygen demand [71].

## Conclusion

Our results indicate that hydrodynamic processes shape the diversity and productivity of cyanobacterial cohorts across Lake Erie, and that these cohorts in turn support the abundance, diversity, and respiration of their heterotrophic consortia. These linkages are taxonomically specific, with the most diverse cohorts (in the central basin) supporting the most diverse and metabolically active consortia. In addition, picocyanobacterial-dominated communities may still support substantial heterotrophic biomass even in the absence of large, visible blooms. Together, these results support a model in which cyanobacterial composition shapes how primary production is transferred into heterotrophic biomass and respiration, offering a mechanistic connection between blooms, microbial diversity, and oxygen demand at a lakewide scale.

While our data suggest a mechanistic link between cyanobacterial productivity and microbial metabolism, additional studies are needed to confirm these processes. Basin-wide measurements of cyanobacterial cohort abundance alongside oxygen demand are needed to test whether higher respiration in the heterotrophic consortia is balanced by greater photosynthetic oxygen production. In addition, interannual studies of central basin microbial communities would allow us to confirm the cohorts and consortia we present here are taxonomically consistent and reoccur annually. Finally, functional measurements, including metatranscriptomics and stable isotope probing, would provide insight to the specific interactions which structure the consortia we observe and their biogeochemical implications.

## ACKNOWLEDGMENTS

We thank Eric Osantowski, Anne Scofield, Ben Alsip, and other staff in the Great Lakes National Program Office of the US EPA and the crew of the *R/V Lake Guardian* for supporting sample collection and metadata. We thank Brayan Vilanova-Cuevas and Elijah Jones for support in May field work. We thank Mark Rowe for coding advice in analyzing NOAA data. We thank Linda Cote and the BRC Genomics Facility (RRID:SCR_021727) at the Cornell Institute of Biotechnology for sequencing experiments. This work was supported by a NOAA Margaret A. Davidson Fellowship to A.P. (NA24NOSX420C0016) and by a grant from the Affinito-Stewart & President’s Council of Cornell Women, as well as Cornell University start-up funds to M.L.S. Toxin analysis was supported by the Lake Erie Center for Freshwater and Human Health under NIH National Institute of Environmental Health Sciences (2P01ES028939-06) and National Science Foundation (OCE-1840715) to G.L.B. Thank you to the anonymous reviewers for their valuable feedback.

## DATA, METADATA, AND CODE AVAILABILITY

All raw and processed data for this project are publicly available. The main GitHub repository for this project is available at https://github.com/MarschmiLab/AAM_ERI_2026, which includes all processed data and code. The raw compressed 16S rRNA gene sequencing fastq files are available in the NCBI Sequence Read Archive under the BioProject #PRJNA1481647. All flow cytometry data are available from Zenodo (https://doi.org/10.5281/zenodo.20836188).

