## Supplemental Methods and Figures for "Cyanobacterial cohorts structure the diversity, abundance, and metabolism of heterotrophic bacteria in Lake Erie"

### FULL MATERIALS AND METHODS

#### Sample collection

Samples were collected in 2024 in collaboration with the US Environmental Protection Agency (EPA) aboard the *R/V Lake Guardian* as part of the Lower Food Web Cooperative Science and Monitoring Initiative (CSMI) surveys May 20-23 and September 13-16, and as part of the annual summer survey August 9-14 (Fig. 1). A total of 33 stations were visited at each cruise; complete sampling dates and locations can be found in Table S1. At all stations, depth profiles were collected via a conductivity, temperature, and depth (CTD) meter equipped with additional sensors for dissolved oxygen, fluorescence, and turbidity.

Depth-discrete water samples for environmental data were collected using an attached Niskin-bottle rosette, with "surface" samples taken at 2 m deep; when stations were sufficiently deep to observe a thermocline, additional "bottom" samples were collected 2 m above the lakebed. Chemical parameters and chlorophyll-a concentrations were analyzed by the US EPA's Great Lakes National Program Office using standard protocols [1].

Microbial sampling was carried out at as many stations as possible, given time constraints of shipboard sampling (Fig. 1). At each station, an integrated epilimnetic sample was generated by combining equal volumes of water from 1 m, 5 m, and 10 m depths and homogenizing by gentle mixing, hereafter referred as just "surface" samples. This sampling approach was used because bloom-forming cyanobacteria like *Microcystis sp.* can be heterogeneous within the epilimnion due to the presence of intracellular gas vesicles that reduce the density of the cells and allow them to modify their buoyancy vertically in the water column [2]. For amplicon sequencing, 2 L of water was filtered through a 0.22 $\mu$m polyethersulfone (PES) filter (MilliporeSigma) and then flash frozen in liquid $N_2$ before storage at -80°C. Filtration lines were sterilized with bleach and MilliQ water between samples. For particulate toxin analysis, 2.5 L of water was filtered through 0.7$\mu$m GF/F glass fiber filters (Whatman) and samples were stored at -20°C. Between toxin samples, filteration lines were rinsed only with MilliQ water. At the end of each cruise, field negative controls for both DNA and toxin filters were produced by running

2L of MilliQ onto either 0.22 $\mu$m PES or 0.7$\mu$m GF/F filters after standard cleaning procedures.

#### Cyanotoxin Quantification

Filters for cyanotoxin analysis were transferred to SUNY-ESF for analysis. They were extracted in 50% acidified methanol using ultrasound, centrifuged at 14,000 x g, and filtered through a 0.22 $\mu$m nyloin filter prior to storage at -20°C. Microcystins (17 congeners) were analyzed by HPLC-coupled with single quadrupole mass spectrometry and quantified against a standard curve of microcystin LR or RR as outlined in Boyer [3]. Anatoxin-a, homo-anatoxin-a, cylindrospermopsin, and deoxy-cylindrospermopsin concentrations were determined using LC-MS/MS as described in Smith et al. [4]. The method detection limits for all toxins was <0.01 $\mu$g/L.

#### Cell quantification via flow cytometry

Flow cytometry was used to quantify three microbial properties: (1) total microbial cell abundances, (2) phototrophic cells based on chlorophyll-a autofluorescence, and (3) metabolically active cells using RedoxSensor Green (RSG) labeling.

Samples for flow cytometry were collected from integrated eplimnetic samples prefiltered through sterile 200$\mu$m and 20 $\mu$m Nitex mesh (Wildco) to prevent clogging. All counts were performed using an Attune Nxt flow cytometer (Thermofisher) equipped with a small-particle filter. For total cell counts and phototrophic cell measurements, 1.5 mL of sample was fixed with 1.5$\mu$L of 25% glutaraldehyde for 10 minutes at room temperature. For metabolic labeling with RSG, which was only collected in September, 1 mL of prefiltered sample was first incubated with 1 $\mu$L of 1mM RSG (Thermofisher) for 30 minutes in the dark at room temperature, after which 1$\mu$L of 25% glutaraldehyde was added and samples were incubated for an additional 10 minutes. All samples were collected in biological duplicates, flash frozen in liquid $N_2$, and stored at -80°C.

Total cells were quantified based on DNA staining with SYBR Green I [5]. Fixed microbial samples were thawed for 30 minutes at 37 °C, diluted 10x in sterile PBS, and stained in technical duplicates with SYBR Green I dye (Bio-Rad) at a final concentration of 1x at 37 °C for 20 minutes in the dark. Total cells were counted using BL1-H fluorescence (excitation: 488 nm, emission: 530/30), a sample volume of 50 $\mu$L, and a 25 $\mu$L/min flow rate. DNA+ cells were defined using a BL1-H gate (Fig. S1).

Phototrophic cells were quantified based on chlorophyll-a autofluorescence [6]. Thawed samples were diluted 2x in sterile PBS and Chl+ cells were counted in technical duplicates using BL3-H fluorescence (excitation: 488nm, emission: 695/40), with a sample volume of 150 $\mu$L, and a 25 $\mu$L/min flow rate. Chl+ cells were defined using BL3-H fluorescence gate (Fig. S1). Cell size was estimated using the forward scatter (FSC-H) after creating a standard curve using a Flow Cytometry Size Calibration Kit (Thermofisher; F13838), with a 5$\mu$m cutoff for "Large" cells (Fig. S1).

Metabolically active cells were quantified using RedoxSensor Green (RSG), which is reduced by intracellular oxidoreductases associated with cellular respiration, producing fluorescence proportional to metabolic activity [7]. Thawed samples were diluted 10x in sterile PBS; half of the sample was left stained only with RSG (RSG-only), whereas the other half was stained in technical duplicates with a DNA stain (SYBR I Green; Bio-Rad) as described above for total cell quantification. Cells were counted using BL1-H fluorescence, a sample volume of 50 $\mu$L, and a 12.5 $\mu$L/min flow rate. Metabolically active cell concentrations were calculated using a BL1-H gate on the SYBR-stained samples, while RSG+ cells were quantified using a separate BL1-H gate on RSG-only samples. RSG+ staining resulted in lower total cell counts compared to DNA-stained samples (median 18.7% decline). When calculating % RSG+ cells, the total cell count within the RSG-stained samples was used as the denominator.

Flow data was analyzed using the R packages flowCore, ggcyto, and custom functions sorted in the flow.gus package. All gates used in analysis are defined within the analysis/Flow_Cytometry scripts in the Github repository associated with this manuscript. Full collection parameters, laser voltages, and thresholds are available within Table S2.

#### DNA Extraction and Metabarcoding

All DNA extractions were carried out using the Qiagen DNeasy PowerWater kit per manufacturer's protocol. Half a filter was used corresponding to 1 L of water extracted, and an extraction negative was produced for each kit using a blank filter.

The V4–V5 hypervariable region was amplified using the universal primers 515F (5-GTGYCAGCMGCCGCGGTAA) and 926R (5-CCGYCAATTYMTTTRAGTTT) as described in Yeh et al. [8] and Needham et al. [9]. Triplicate 25 $\mu$L polymerase chain reactions were performed with KAPA HiFi 2x MasterMix (Roche) with primers at an end concentration of 3$\mu$M and 5 ng of sample DNA. In total, 80 microbial samples were sequenced along with negative controls (four field blanks, two extraction blanks, one PCR blank, and one indexing blank) and a defined microbial community (ZymoBIOMICs Microbial Community DNA Standard) used as a positive control to assess amplification error rates. Libraries were sequenced at the Cornell Biotechnology Resources Center using a 2 x 300 bp paired-end XLEAP kit on an Illumina NextSeq 2000, yielding 62,808,466 reads with a median of 879,695 reads/sample (excluding blanks).

Sequencing data were processed to generate amplicon sequence variants (ASVs) from both prokaryotic and eukaryotic reads using scripts modified from McNichol et al. [10]. Primers were removed with Cutadapt and reads were classified as either prokaryotic (16S) or eukaryotic (18S) using bbsplit [11, 12]. Prokaryotic sequences were denoised and ASVs inferred using a standard DADA2 workflow, with minor adjustments for binned quality scores [13]. Eukaryotic sequences are not used here, but are publicly available within the associated Github repository.

Prokaryotic ASVs were taxonomically classified using the Greengenes2 database (v2024.09) [14]. Mitochondrial and chloroplast ASVs were removed, leaving 17,407 inferred ASVs. The mock community contained eight erroneous ASVs, likely resulting from sequencing errors, representing 0.5% of the sequencing reads in the mock community. Two additional ASVs were removed because they were present in the negative controls or appeared to originate from mock community contamination. To reduce potential erroneous ASVs, a minimum observation threshold of eight reads per ASV was applied, leaving 10,774 ASVs but retaining 99.9% of all sequencing reads.

Phylogenetic trees were constructed for both eukaryotic and prokaryotic datasets using MAFFT for sequence alignment and FastTree for tree inference under a generalized time-reversible model [15, 16]. Four ASVs producing anomalously long branches in the prokaryotic tree were removed. Archaea were poorly represented (three ASVs); given previous documentation of archaeal taxa in Lake Erie, this scarcity likely reflects low amplification efficiency for Arhcaea with these primers despite positive *in silico* predictions [10, 17]. Sampling depth for prokaryotes was assessed using rarefaction, and a minimum of 10,000 reads per sample was applied. One sample (Stn. 937 in September) exceeded this threshold (14,676) but exhibited anomalously low richness and cell counts, suggesting possible residual bleach contamination in the sampling bottle, and was therefore removed. Of the original 80 samples sequenced, nine were excluded from final analyses due to insufficient sequencing depth or quality (Table S3).

#### Ecological analyses

Alpha-diversity (richness) was estimated after rarefying samples to the minimum sequencing read depth (11,901 reads) using custom scripts in the fun.gus package [18].

Beta-diversity was calculated using Bray-Curtis dissimilarity based on absolute abundances. ASV absolute abundances were calculated by first converting all ASVs to their relative abundance per sample. Each ASV's absolute abundance in a sample was then calculated by multiplying its relative abundance by the total cell count of that sample as determined by flow cytometry [19]. Because flow cytometry counts were generated from the <20 $\mu$m fraction whereas sequencing was performed without any prefiltration, colonial cyanobacteria were likely underestimated during flow cytometry and compared to their total relative abundance in sequencing reads; as a result, most samples have a greater number of total predicted cyanobacterial counts (sequencing) compared to total phototroph counts (flow cytometry, mean difference: 173,313 cells/mL). Despite this discrepancy, the two measurements were strongly correlated (Spearman's $R = 0.82$, $p < 0.0001$). Cyanobacterial spatial groups were defined using UPGMA clustering, cutting the tree at 7 groups to maximize clustering by both month and basin.

Co-occurrence network analyses were performed using functions from the igraph and tidygraph packages [20, 21]. For cyanobacteria–cyanobacteria co-occurrence, the 100 most abundant cyanobacterial ASVs were selected by summing absolute abundances across all samples in August and September, requiring a prevalence of at least 25%. Pairwise associations were assessed using Pearson correlations, and significant correlations (edges) were defined as $r \geq 0.8$ and $p \leq 0.01$ (uncorrected). These edges were used to construct a weighted, undirected network that maximized modularity (Fig. S2). For the cyanobacteria–heterotroph co-occurrence network, the 1000 most abundant ASVs in August and September with a prevalence of at least 25% were first selected (excluding the 100 most abundance cyanobacterial ASVs). Because the initial network was dominated by cyanobacteria–cyanobacteria relationships (Fig. S3), cyanobacterial ASVs were removed and a minimum absolute abundance of $> 50{,}000$ cells/ml was applied, leaving 344 heterotrophic ASVs. Pearson correlations were then calculated between each of the 100 cyanobacterial ASVs ("hubs" and abundant non-cyanobacterial ASVs ("nodes"). Significant correlations (edges) were defined as $r \geq 0.85$ and $p < 0.01$ (uncorrected) and used to construct a weighted, undirected bipartite network. For both networks, clusters were identified using the Givan-Newman algorithm implemented in the `cluster_edge_betweenness` function, and networks were visualized using the Fruchterman-Rheingold layout.

General data manipulations relied on the tidyverse and phyloseq packages using R v4.3.3 [22, 23, 24]. Other statistical tests (including Two-Sample Wilcoxon Tests and Spearman Correlations) were performed using functions from base R, rstatix, or vegan packages [25, 26]. Plots were produced using functions from the ggplot2, ggpubr, and ggdendro packages [23, 27, 28].

### Supplemental Figures

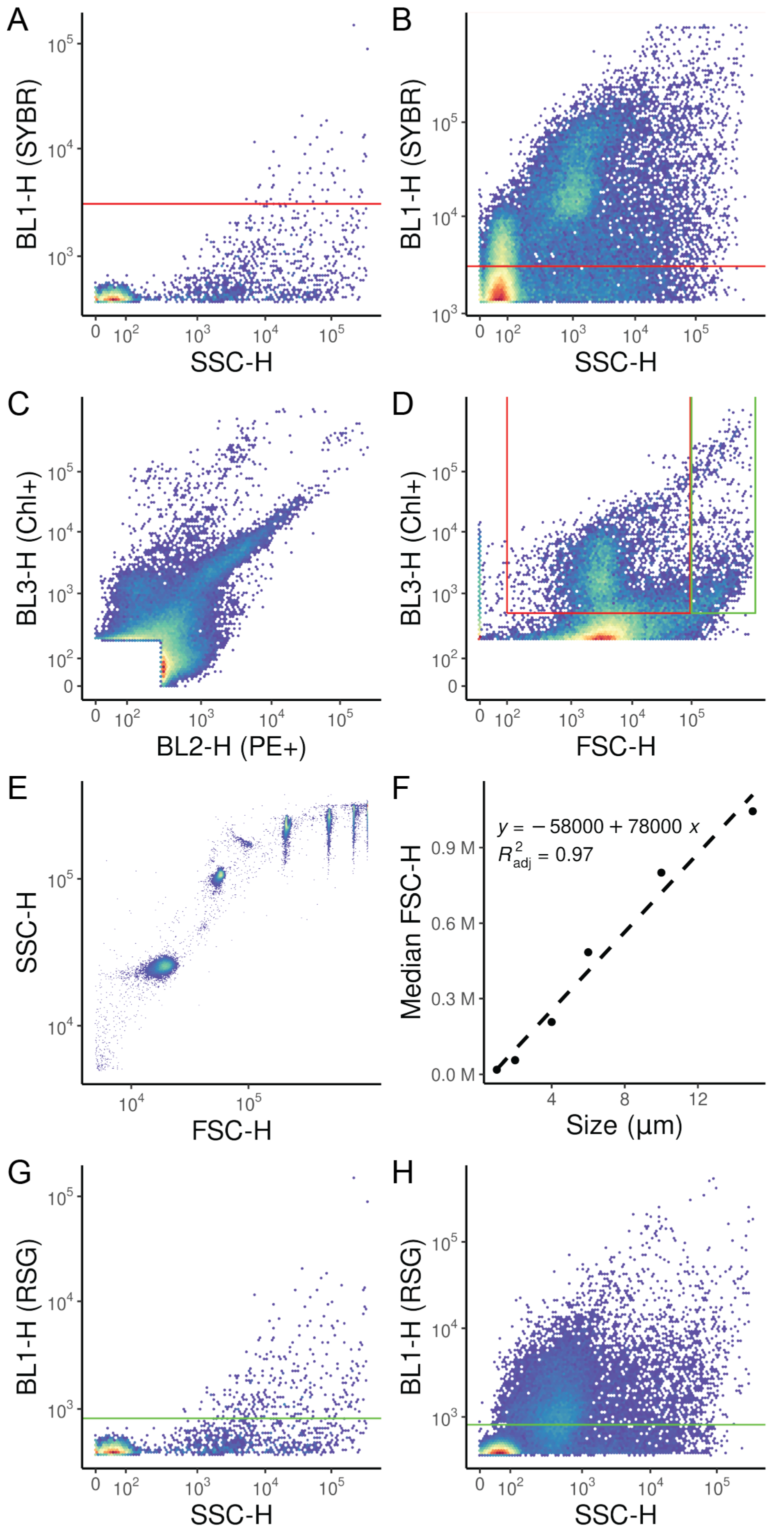


**FIG** S1 *Flow cytometry gating and size estimation* All panels besides E-F are from Station 58 in September. (A) Unstained sample under cell counting conditions, with the SYBR+ (cell) gate in red, demonstrating few false positives (B) The same sample stained with SYBR-green, with the SYBR+ gate in red. (C) Example thresholding using BL2-H and BL3-H to detect chlorophyll or phycoerythrin in an unstained sample. (D) Example gating for "small" Chl+ phototrophs (red gate) and "big" Chl+ phototrophs (green gate). (E) Size selection beads demonstrating linear FSC-H response. (F) Standard curve using the median FSC-H values for each standardized bead size. (G) Unstained cells under RSG-counting conditions, with the RSG+ gate in green. (H) The same sample with RSG+ positive staining, with the RSG+ gate in green.

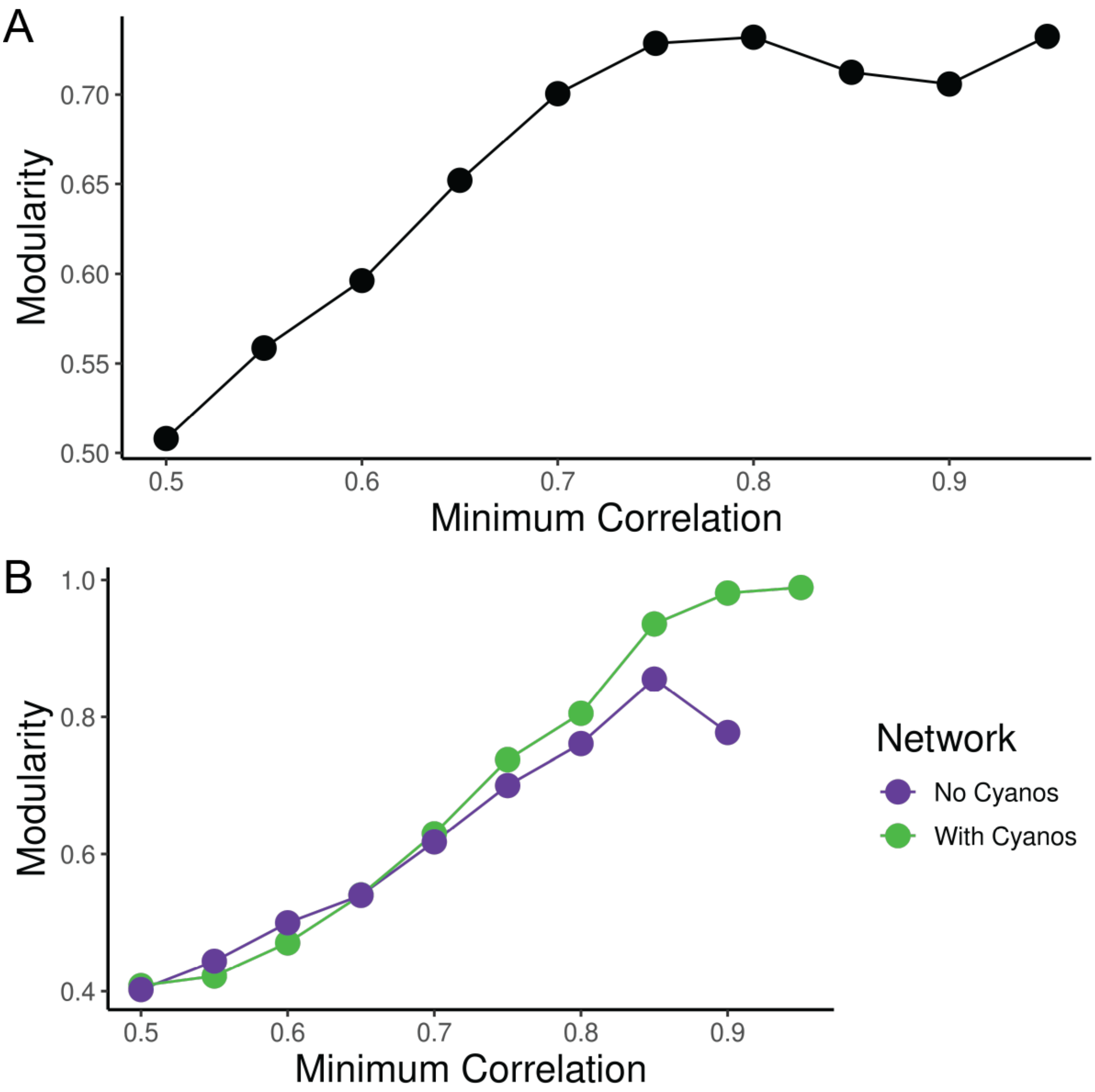


**FIG** S2 *Network modularity across significant correlation thresholds.* Graphs were constructed from correlation matrices, defining significant edges using $p < 0.01$ and increasingly high Pearson correlations (x-axis). Networks were clustered using the Givan-Newman algorithm, and modularity was measured on the clustered graphs. These analyses were run for the 100 most abundant cyanobacterial ASVs (A, Fig. 6) or the 100 most abundant cyanobacterial ASVs versus the *other* 344 abundant and prevalent heterotrophic ASVs (B, Fig. 8), excluding other cyanobacterial ASVs.

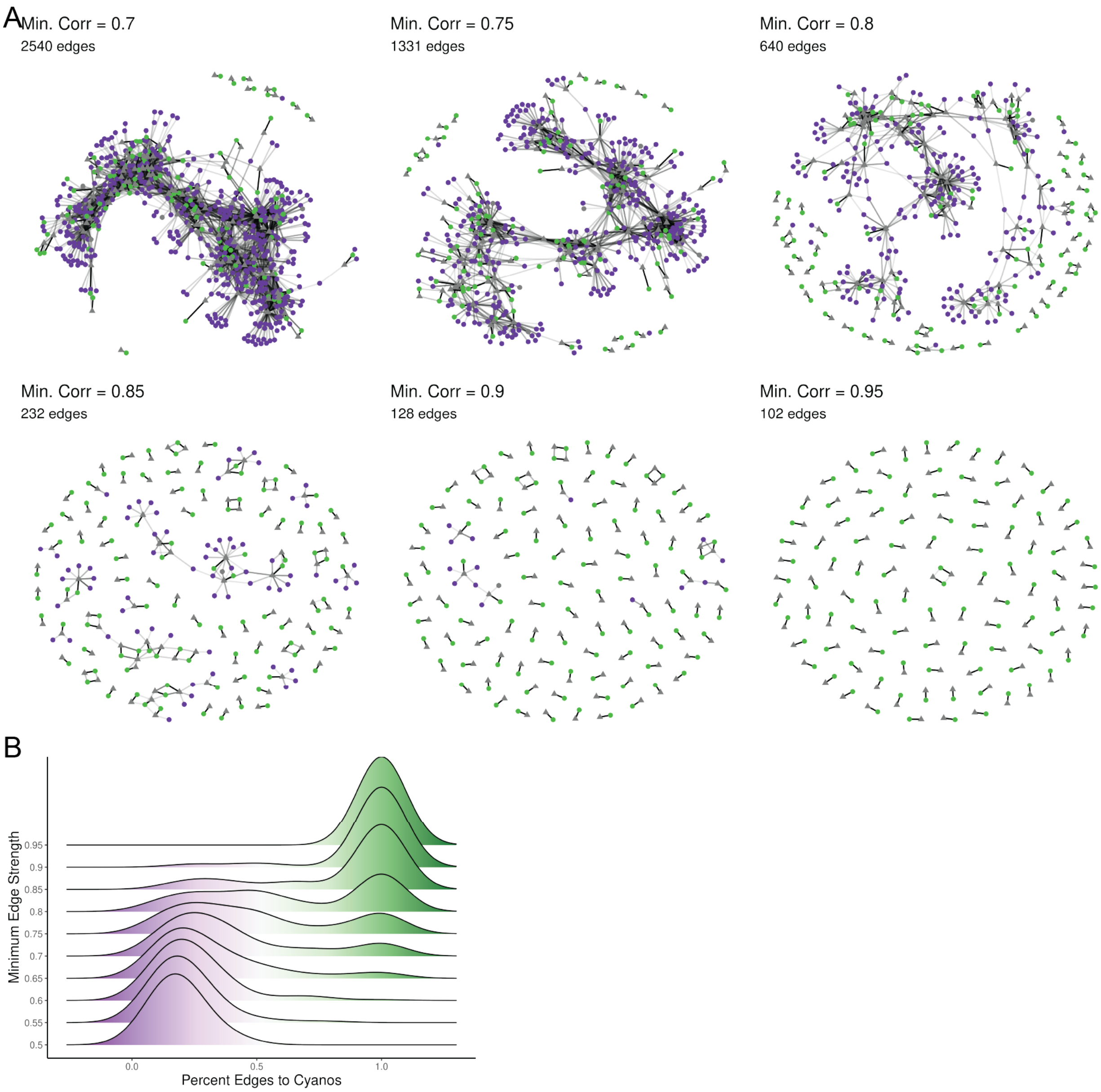


**FIG** S3 *Cyanobacteria form the strongest cooccurence relationships with each other.* (A) Cooccurrence networks across increasingly high correlation thresholds for edges. The 100 most abundant cyanobacterial "hubs" are shown as gray triangles, while connected nodes are either green (for other cyanobacteria) or purple (for other non-cyanobacteria). (B) Distributions of the percent of edges which are formed between two cyanobacterial ASVs as the minimum correlation threshold for an edge is increased.

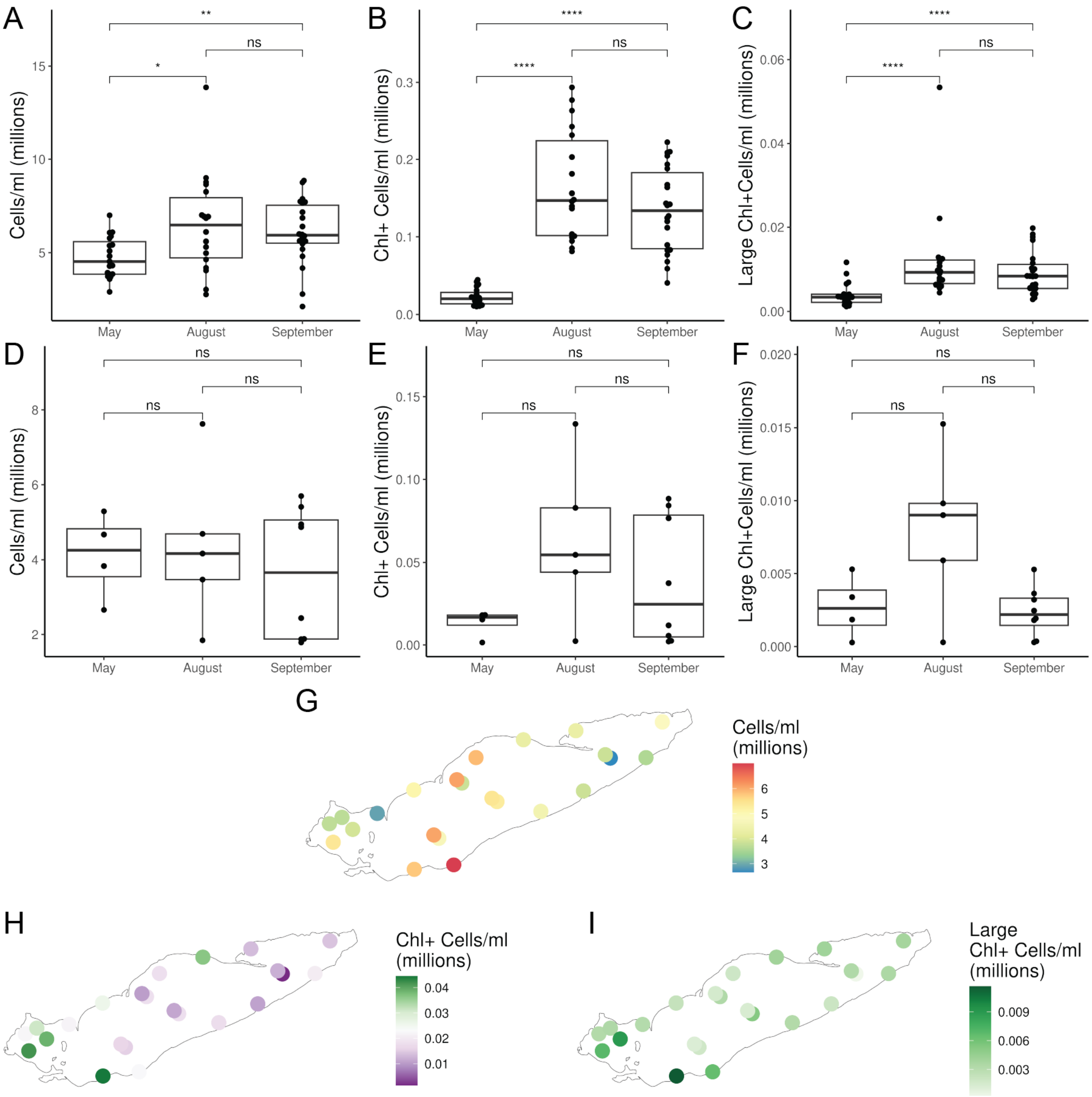


**FIG** S4 *Cell counts and phototrophs in bottom and May samples.* (A/D) Total bacterial cell abundance, (B/E) Chlorophyll-a + cells, and (C/F) Large (>5$\mu$m) chlorophyll-a + cells were measured via flow cytometry after prefiltering all samples to <20$\mu$m. (A-C) represent surface samples, while (D-F) represent bottom samples. Brackets represent results of Two-Sample Wilcoxon tests with Holm-Bonferroni adjustment for multiple comparisons (ns = p > 0.05, * = p < 0.05, ** = p < 0.01, *** = p < 0.001, **** = p < 0.0001). (G) Total bacterial cell abundance, (H) Chlorophyll-a + cells, and (I) Large (>5$\mu$m) chlorophyll-a + cells in May. Points represent surface samples, with bottom samples for those stations plotted offset underneath.

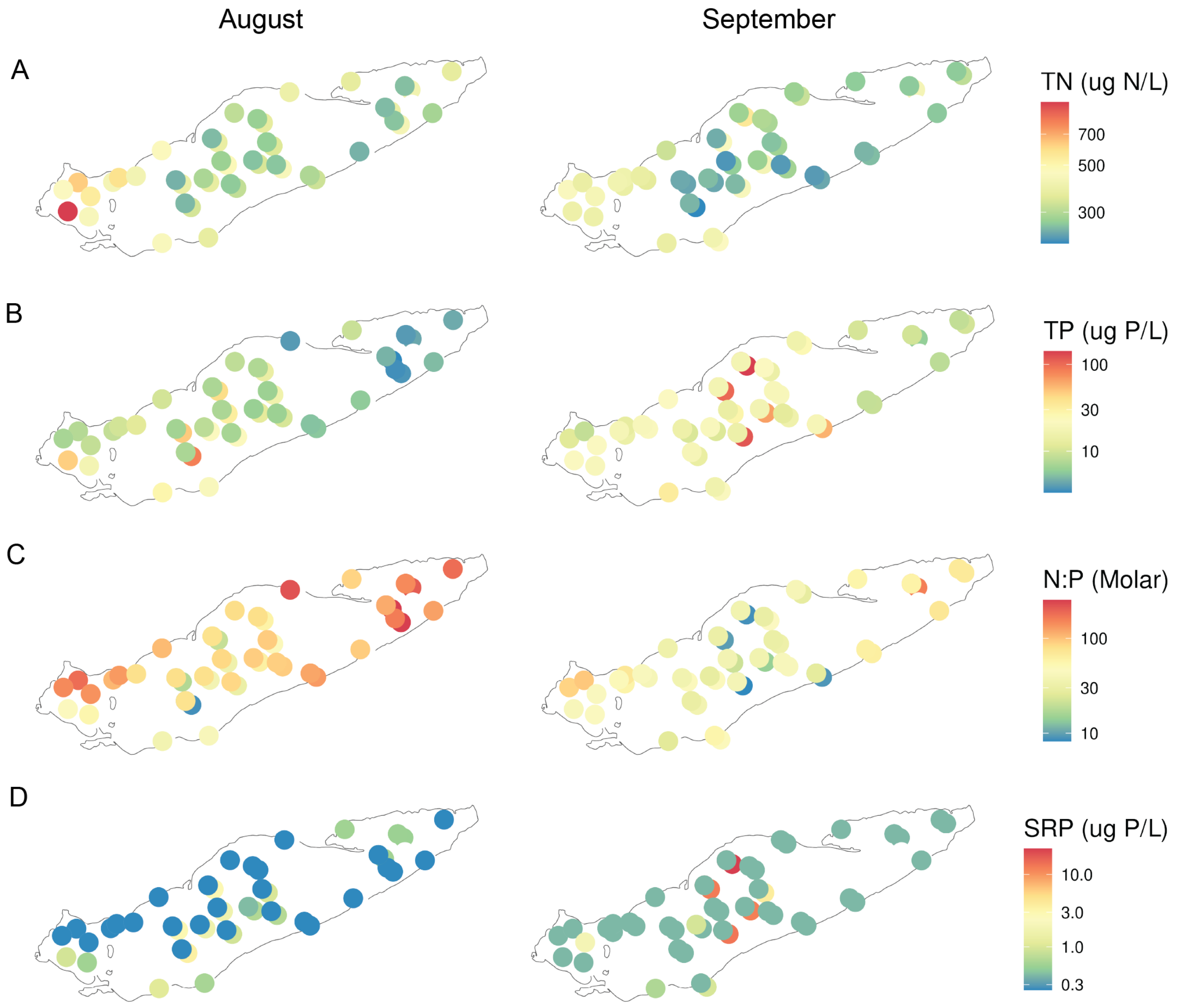


182

**FIG** S5 *Nutrient measurements across Lake Erie* Data is drawn from EPA GLNPO chemical analyses. Bottom points are shown offset beneath surface points. Surface corresponds to the average of any samples taken < 10 m deep, while bottom samples were taken 2 m above the lake bed. For some nearshore stations in August with depths < 10 m, surface samples represent integrated water column samples. Note that nutrient colors are log-scaled.

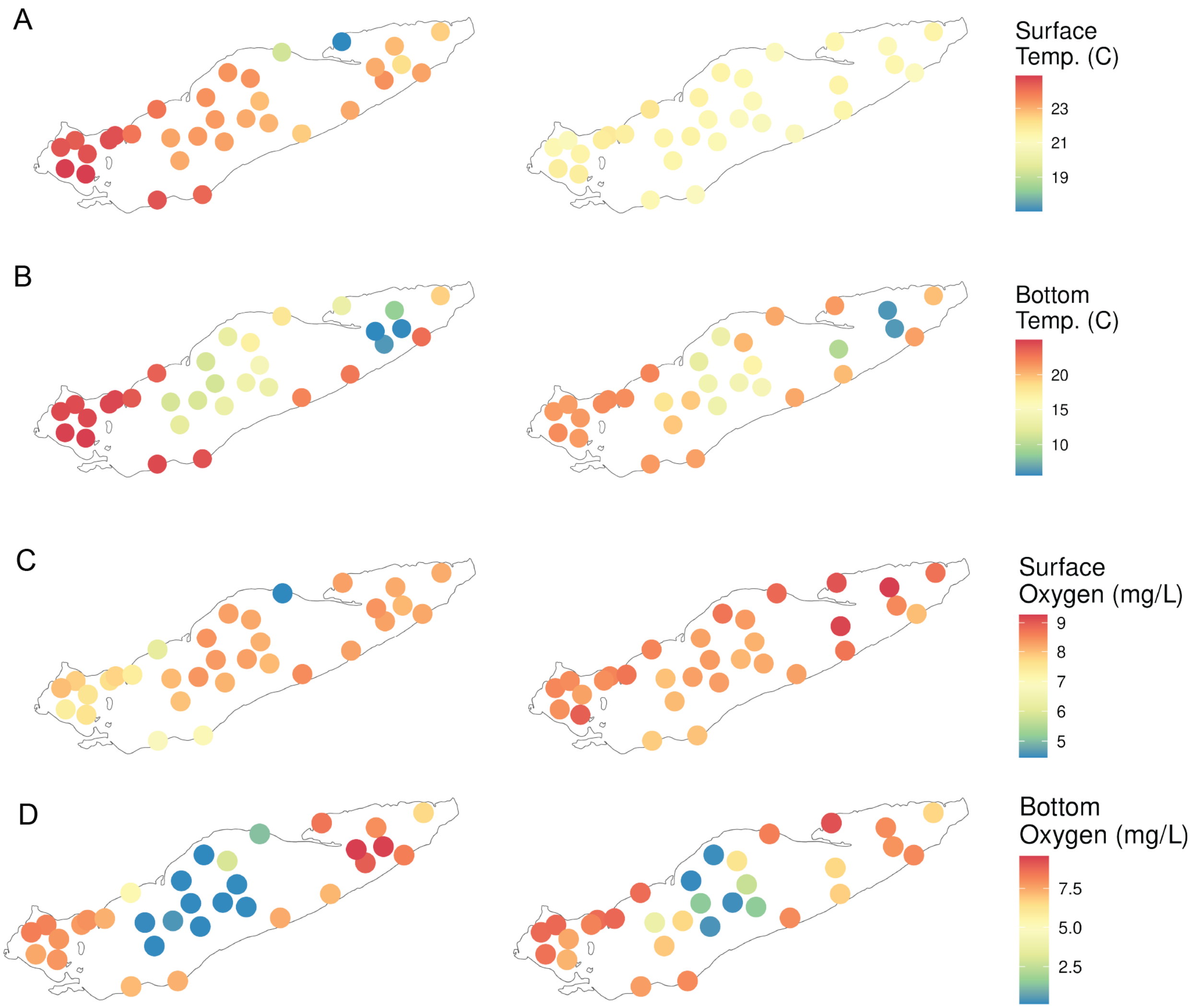


**FIG** S6 *Temperature and oxygen measurements across Lake Erie* Data is drawn from CTD casts. Bottom points are shown offset beneath surface points. Surface measures represent the average reads from 1.5 m - 10 m deep, to match integrated microbial sampling. Bottom samples represent the average of the deepest 1 m of readings, typically 1 m - 2 m above the lake bed.

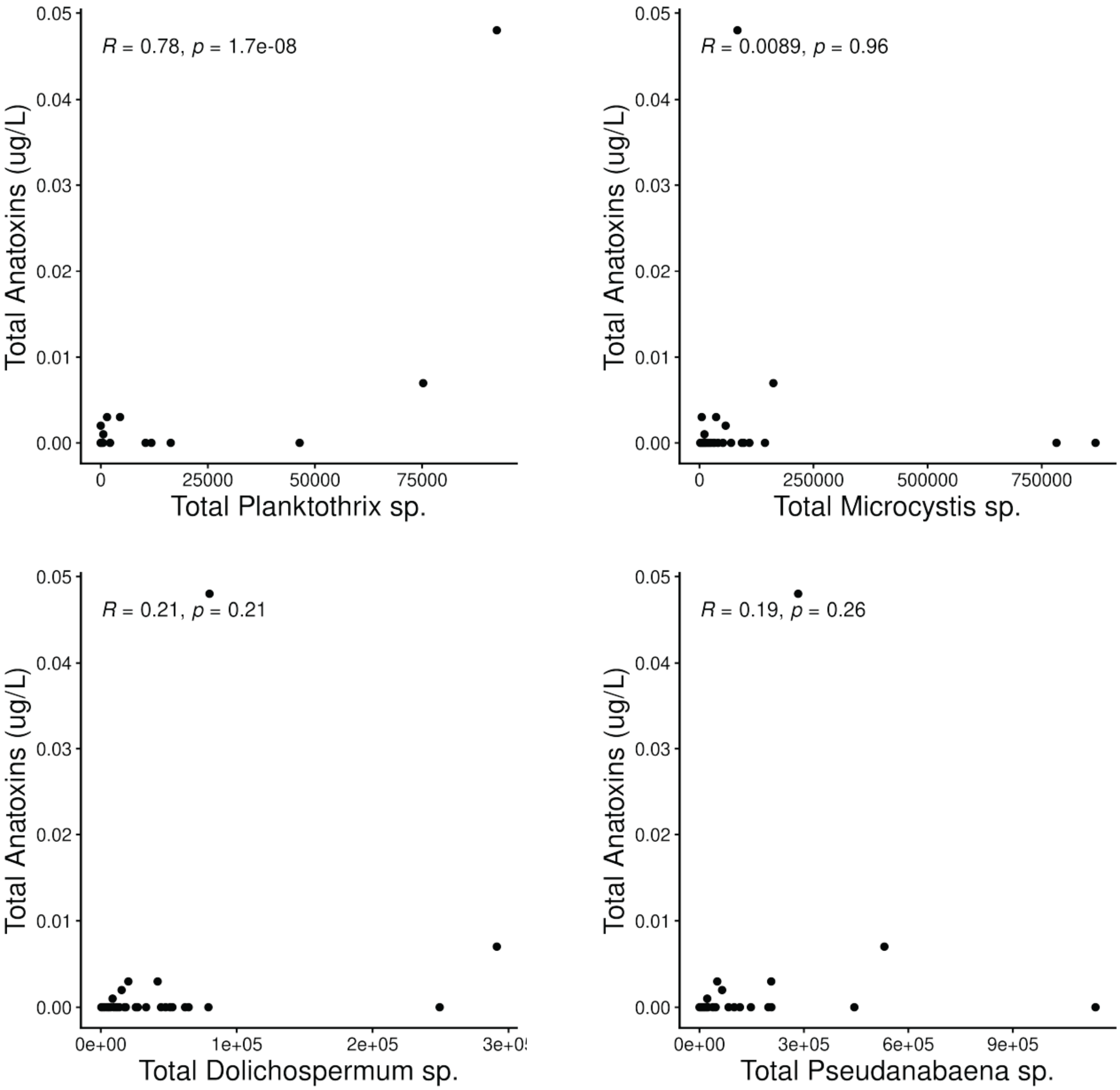


**FIG** S7 *Non-significant correlations between cyanobacterial genera and anatoxins.* Estimated absolute abundance of four cyanoabacteria is shown in each panel, versus the total anatoxins within each sample. R corresponds to Spearman's Rank Correlation.

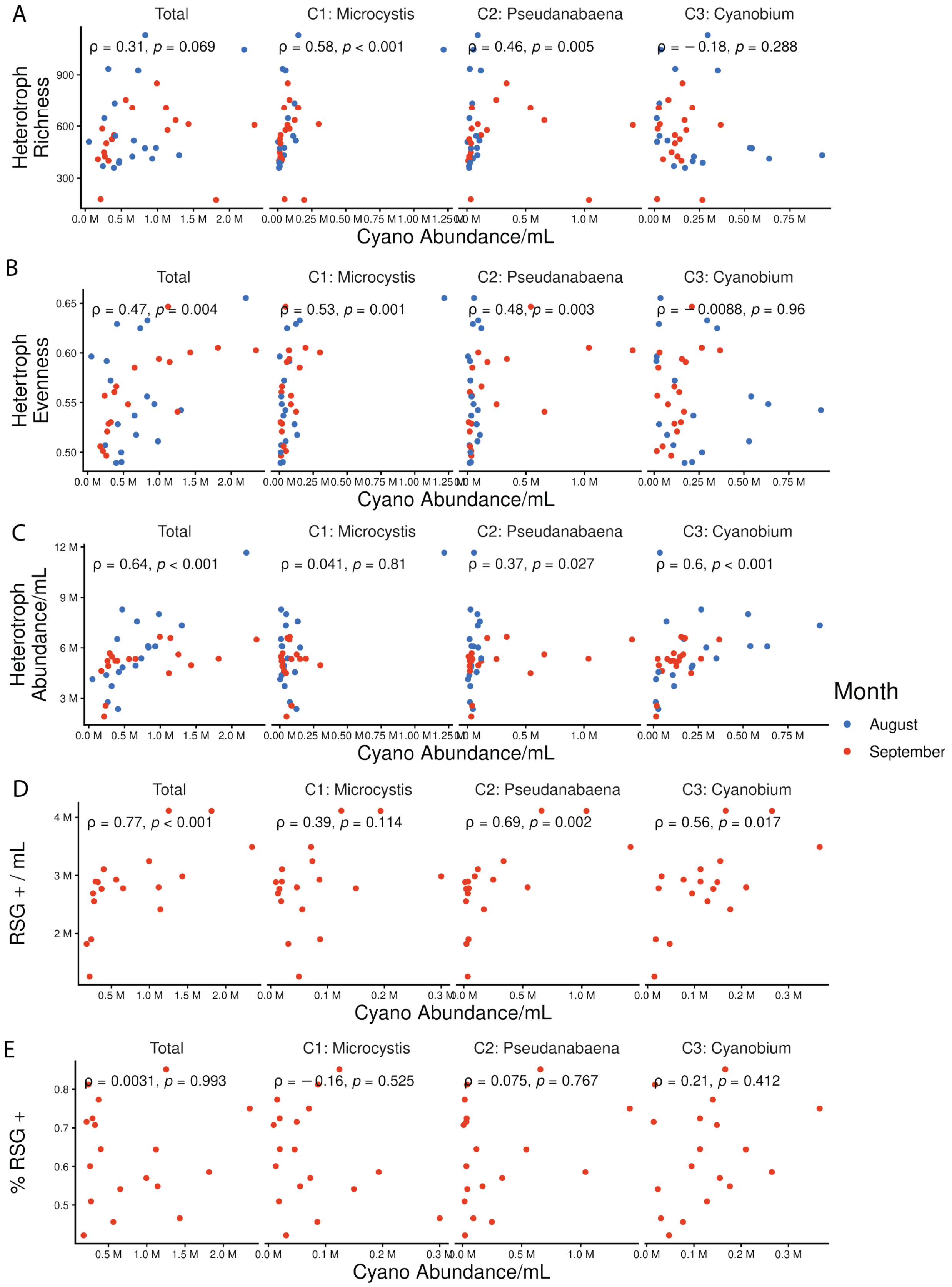
A
B
C
D
E
Total
C1: Microcystis
C2: Pseudanabaena
C3: Cyanobium
Heterotroph Richness
ρ = 0.31, p = 0.069
ρ = 0.58, p < 0.001
ρ = 0.46, p = 0.005
ρ = −0.18, p = 0.288
Hetertroph Evenness
ρ = 0.47, p = 0.004
ρ = 0.53, p = 0.001
ρ = 0.48, p = 0.003
ρ = −0.0088, p = 0.96
Heterotroph Abundance/mL
ρ = 0.64, p < 0.001
ρ = 0.041, p = 0.81
ρ = 0.37, p = 0.027
ρ = 0.6, p < 0.001
RSG + / mL
ρ = 0.77, p < 0.001
ρ = 0.39, p = 0.114
ρ = 0.69, p = 0.002
ρ = 0.56, p = 0.017
% RSG +
ρ = 0.0031, p = 0.993
ρ = −0.16, p = 0.525
ρ = 0.075, p = 0.767
ρ = 0.21, p = 0.412
Cyano Abundance/mL
Month
August
September

**FIG** S8 *Correlations between cyanobacterial cohort abundance and heterotrophic consortia metrics.* All correlations are Spearman's Rank Correlations. Note that p-values aren't corrected between panels; as such, while Spearman's R remains the same between correlations shown here and in Fig. 7, p-values differ, and as such Fig. 7 is more conservative in estimating statistical significant of the correlation.

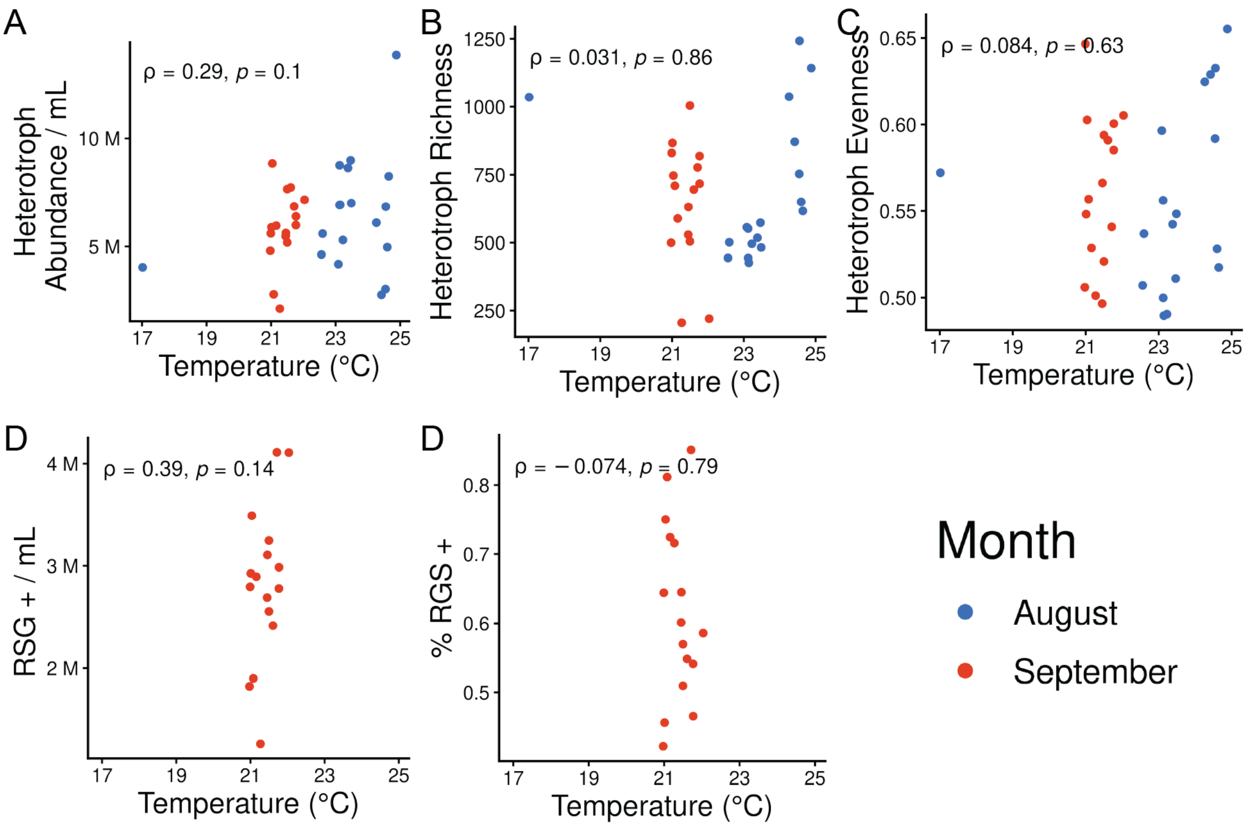


**FIG** S9 *Heterotrophic consortia metrics are not correlated to temperature.* All correlations are Spearman Ranked Correlations (uncorrected across panels, though insignificant regardless).

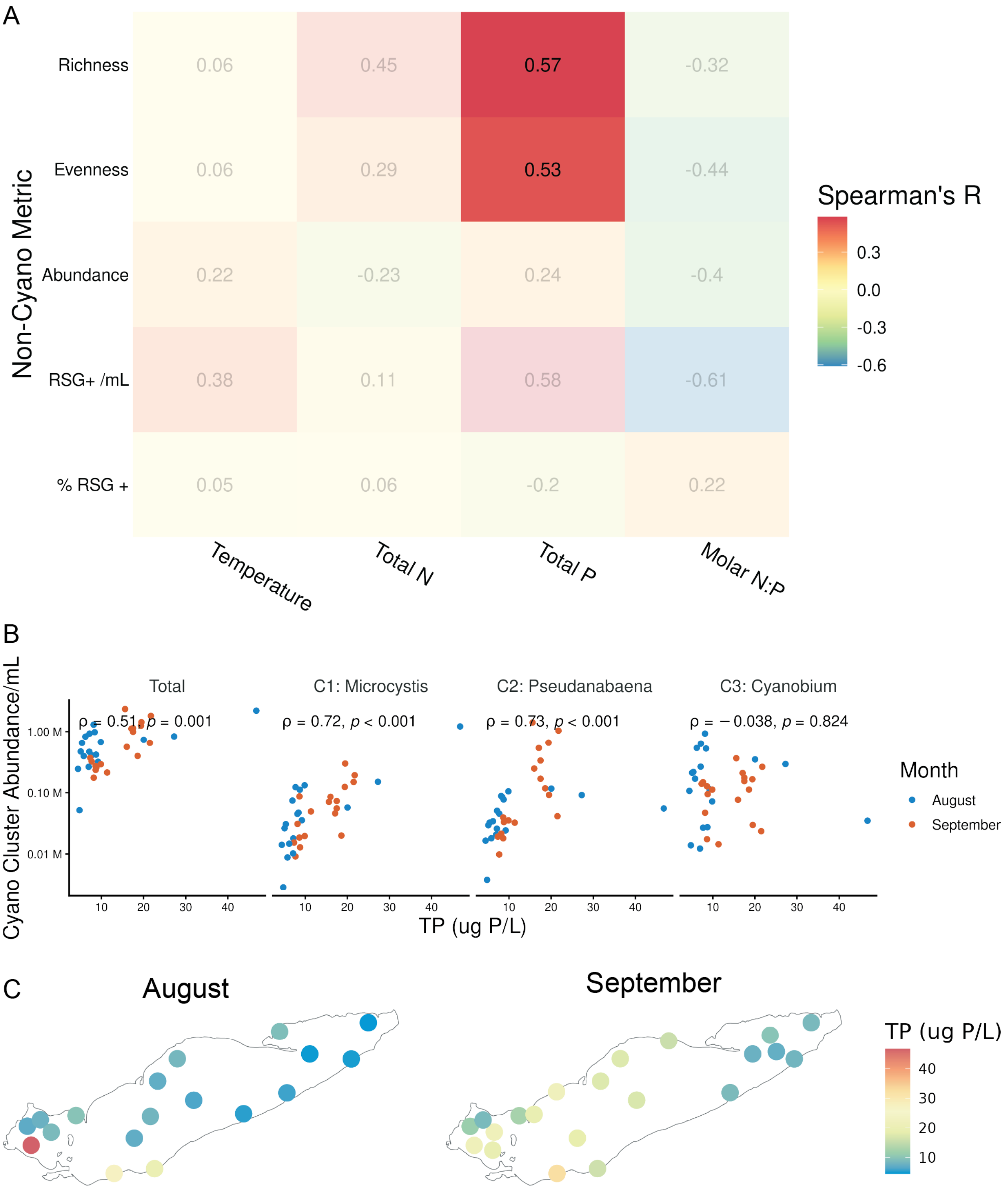


**FIG** S10 *Heterotrophic consortia metrics can not be explained by nutrients* (A) The Spearman Rank Correlation was calculated between nutrient levels and heterotrophic consortia metrics, including total heterotroph richness (minus cyanobacteria ASVs), heterotroph evenness (after removing cyanobacteria), the total cell abundance (minus counts assigned to cyanobacteria), the number of RSG+ (metabolically active) cells, and the % of RSG+ cells in the community. For each comparison, color represents the Spearman Rank correlation coefficient, with the actual correlation provided. Transparent panels were statistically insignificant ($p > 0.05$ with a Holm-Bonferroni adjustment). RSG results are only available for September. (B) Spatial spread of total phosphorus (TP), restricted to surface samples being used in Panel A. (C) Correlations between TP and the sum abundance of cyanobacterial cohorts defined in Fig. 4.

### REFERENCES

[28] **de Vries A, Ripley BD**. 2024. ggdendro: Create dendrograms and tree diagrams using ggplot2.
